# Humanized substitutions of *Vmat1* in mice alter amygdala-dependent behaviors associated with the evolution of anxiety

**DOI:** 10.1101/2021.05.18.444749

**Authors:** Daiki X. Sato, Yukiko U. Inoue, Yuki Morimoto, Takayoshi Inoue, Nahoko Kuga, Takuya Sasaki, Yuji Ikegaya, Kensaku Nomoto, Takefumi Kikusui, Satoko Hattori, Giovanni Sala, Hideo Hagihara, Tsuyoshi Miyakawa, Masakado Kawata

## Abstract

The human vesicular monoamine transporter 1 (*VMAT1*) harbors unique substitutions (Asn136Thr/Ile) that affect monoamine uptake into synaptic vesicles. These substitutions are absent in all known mammals, suggesting their contributions to distinct aspects of human behavior modulated by monoaminergic transmission, such as emotion and cognition. To directly test the impact of these human-specific mutations, we introduced the humanized residues into mouse *Vmat1* via CRISPR/Cas9-mediated genome editing and examined changes at the behavioral, neurophysiological and molecular levels. Behavioral tests revealed reduced anxiety-related traits of *Vmat1*^Ile^ mice, consistent with human studies, and electrophysiological recordings showed altered oscillatory activity in the amygdala under anxiogenic conditions. Transcriptome analyses further identified amygdala-specific changes in the expression of genes involved in neurodevelopment and emotional regulation, which may corroborate the observed phenotypes. This knock-in mouse model hence provides compelling evidence that the mutations affecting monoaminergic signaling and amygdala circuits have contributed to the evolution of human socio-emotional behaviors.

## Introduction

Recent years have seen a surge in the study revealing distinct human brain characteristics at the molecular, cellular, and circuit levels in both cortex and subcortical structures (1–4). Differences in brain monoaminergic signaling are likely to be key factors conferring human attributes, as various monoaminergic pathways are involved in emotional regulation, memory consolidation, cognitive flexibility, and complex social behaviors (5–7). Comparative studies have indeed demonstrated the distinct monoaminergic mechanisms in the human brain (2, 8). Moreover, diversification of monoaminergic mechanisms has been linked to the evolution of social behavior in non-human primates (9, 10). Thus, genetic differentiation of monoaminergic genes may underlie our unique emotional and social characteristics, such as empathy and altruism (2).

In line with these studies, we recently reported that the vesicular monoamine transporter 1 (*VMAT1*; also known as *SLC18A1*) gene has been under positive selection in the human lineage as evidenced by two amino acid substitutions (Glu130Gly and Asn136Thr) not shared by other primates (11). The VMATs transport monoamine neurotransmitters into synaptic vesicles, thereby regulating the kinetics of synaptic transmission. There are two isoforms of VMAT, VMAT1 and 2, with the VMAT2 being highly expressed in the brain and widely studied (12, 13). In contrast, VMAT1 was previously thought to be expressed solely in the peripheral nervous system and chromaffin cells, and its functional importance in the CNS has only recently been implicated (14). Our previous study showed that a new mutation at the 136th amino acid (136Ile) of *VMAT1* has emerged relatively recently in the course of human evolution, and that the human-specific polymorphism Thr136Ile has likely been maintained by balancing selection within non-African populations (11). Fluorometric assays additionally demonstrated that these human-specific residues enhance monoamine neurotransmitter uptake, suggesting the tendency of lower monoamine uptake in early hominids (15). The 136Thr, the dominant variant across populations, is known to be marginally associated with psychiatric diseases, such as bipolar disorder, depression and anxiety, as well as with the neuroticism, a personality trait conferring greater risk of psychiatric disease (16–18). Thus, higher levels of depression and/or anxiety-like behaviors with 136Thr may have been favored in our ancestors, while more recent selection may have led to the emergence of 136Ile after the migration out of Africa and into harsher environments.

Although the evolution of *VMAT1* and its basic functions in the mammalian CNS have been clarified, their contributions to unique human attributes are still unclear. Here we examine the molecular, neurological, and behavioral changes in mice caused by human-specific *VMAT1* mutations. We show that humanized *Vmat1* (*Vmat1*^Ile^) induces amygdala-specific changes in gene expression and neuronal activity, reduces anxiety-like behaviors under anxiogenic conditions and even enhances preference for social novelty. These findings suggest that altered monoaminergic signaling in the amygdala has contributed to the evolution of human emotional and social behaviors, and the present work provides a new experimental strategy for evolutionary studies of human emotions.

## Results

### Conservation of the 136th residue of VMAT1 across non-human vertebrates

To investigate the evolutionary significance of the 136th residue in human VMAT1 (hVMAT1), we first examined the conservation of this residue in a wide range of taxa in vertebrates. Multiple sequence alignments of VMAT1 among 236 vertebrate species revealed the unique evolution of this residue in the human lineage (Fig. 1a). Notably, we confirmed that all species but bicolor damselfish retain asparagine (Asn) at the 136th amino acid, while *Homo sapiens* is the only vertebrate species that carries the unique Thr136Ile (rs1390938) polymorphism.

**Figure 1.**
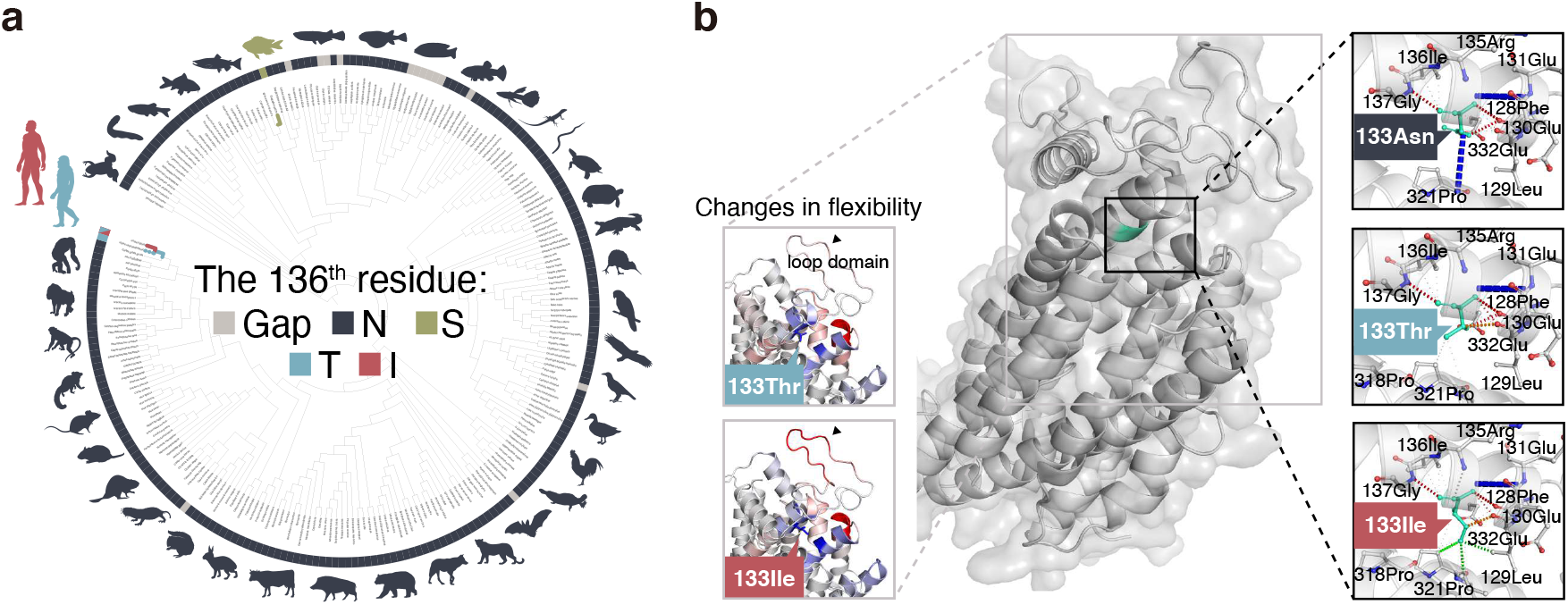
An ultra-conserved residue in VMAT1 exhibits functional variants unique to humans. (a) A phylogenetic tree constructed by multiple sequence alignment of the *VMAT1* gene. Almost all genes across 236 vertebrate species retain asparagine (N, Asn) on the 136th residue, while humans are the only vertebrate species except for bicolor damselfish (shown in green) with a unique polymorphism (T/I, Thr136Ile; rs1390938). Note that the phylogenetic relationship presented here is not necessarily consistent with the known species tree. **(b)** mVMAT1 protein structure predicted by homology modeling and the effects of human-type mutagenesis (corresponding to 133Thr and 133Ile in mVMAT1). (Left) Δ Vibrational entropy energy between WT (133Asn) and mutants. Amino acids are colored according to the vibrational entropy change conferred by the given mutation. Red represents a gain of flexibility and blue represents a rigidification of the structure. The 133Ile mutation leads to increased flexibility of the first luminal loop, a receptor-like domain affecting the affinity of ligands. (Right) When introduced *in silico*, 133Ile but not 13Thr exhibits hydrophobic interactions (shown by green dotted line) with surrounding sites, which likely influence the folding and/or stability of mVMAT1 protein. Blue, red, and orange dotted lines represent amide bonds, hydrogen bonds, and weak van der Waals interactions, respectively.

### Predicted effects of humanized residues on mouse VMAT1 structure and function

*In silico* predictions of wild type mouse VMAT1 (mVMAT1) protein structure revealed that like the hVMAT1 residue 136, the corresponding residue in mVMAT1 (133Asn) was located between the first luminal loop and the second transmembrane domain (Fig. 1b), suggesting a similar function. Given the evolutionary conservation across vertebrates and the difference in chemical properties between amino acids, the humanized mutations to mVMAT1, especially 133Asn to 133Ile, were predicted to have a significant impact on protein function (Table 1). Furthermore, the Asn133Ile mutation was also predicted to stabilize the protein structure as evidenced by greater folding free energy (ΔΔG: 1.106 kcal/mol; Table 1) and to increase the flexibility of the loop domain (Fig. 1b), which possibly increases the efficiency of monoamine uptake.

**Table 1.** In silico prediction of mVMAT1 tolerance to the humanized mutations Asn133Thr and Asn133Ile. Text in bold indicates significant effects of the substitutions as predicted by Provean score < –2.5 or SIFT score < 0.05. The Provean term “Deleterious” and SIFT term “Damaging” refer to significant effects on protein function but do not necessarily indicate that they deteriorate protein function. All three methods predict that 133Ile has greater effects than 133Thr on mVMAT1 function.

| Ref. | Alt. | Provean |  | SIFT |  | DynaMut |  |
| --- | --- | --- | --- | --- | --- | --- | --- |
| | | Score | Prediction | Score | Prediction | $\Delta\Delta G$<br>(kcal/mol) | Prediction |
| Asn | Thr | **-3.27** | **Deleterious** | 0.243 | Tolerated | -0.062 | Destabilizing |
| Asn | Ile | **-6.55** | **Deleterious** | **0.014** | **Damaging** | 1.106 | Stabilizing |

### Generation of *Vmat1*-humanized mouse models by CRISPR/Cas9 genome editing

In order to replace the 133Asn of mouse Vmat1 with Thr or Ile, we employed the CRISPR/Cas9-mediated genome editing strategy (Fig. 2a). We selected a guide RNA targeting the 133rd Asn codon in *Vmat1* exon 4 by using the web-based CRISPR design tool to minimize off-target cleavage risk (Supplementary Table S1 and 2, see Methods for details). By electroporating CRISPR components (*Vmat1* exon 4 crRNA, tracrRNA, and Cas9 nuclease) along with the ssDNA donor containing the desired substitutions into C57BL/6J fertilized eggs (Fig 2a, Supplementary Fig S1a), we successfully generated 133Thr founder No.1 and 133Ile founder No.10 (Supplementary Fig. S1b, c and d). After confirming the absence of off-target cleavages (Supplementary Fig. S2), we crossed 133Thr founder No.1 and 133Ile founder No.10 with wild type (WT) C57BL/6J mice to obtain heterozygous F1 generations, which were again verified to possess the designed substitutions by Sanger sequencing. The homozygous mice were further crossed after the F5 generations to obtain three humanized genotypes, Thr/Thr, Thr/Ile, and Ile/Ile, in addition to WT, for behavioral, neurophysiological, and transcriptomic analyses (Supplementary Fig. S3, see Methods).

**Figure 2.**
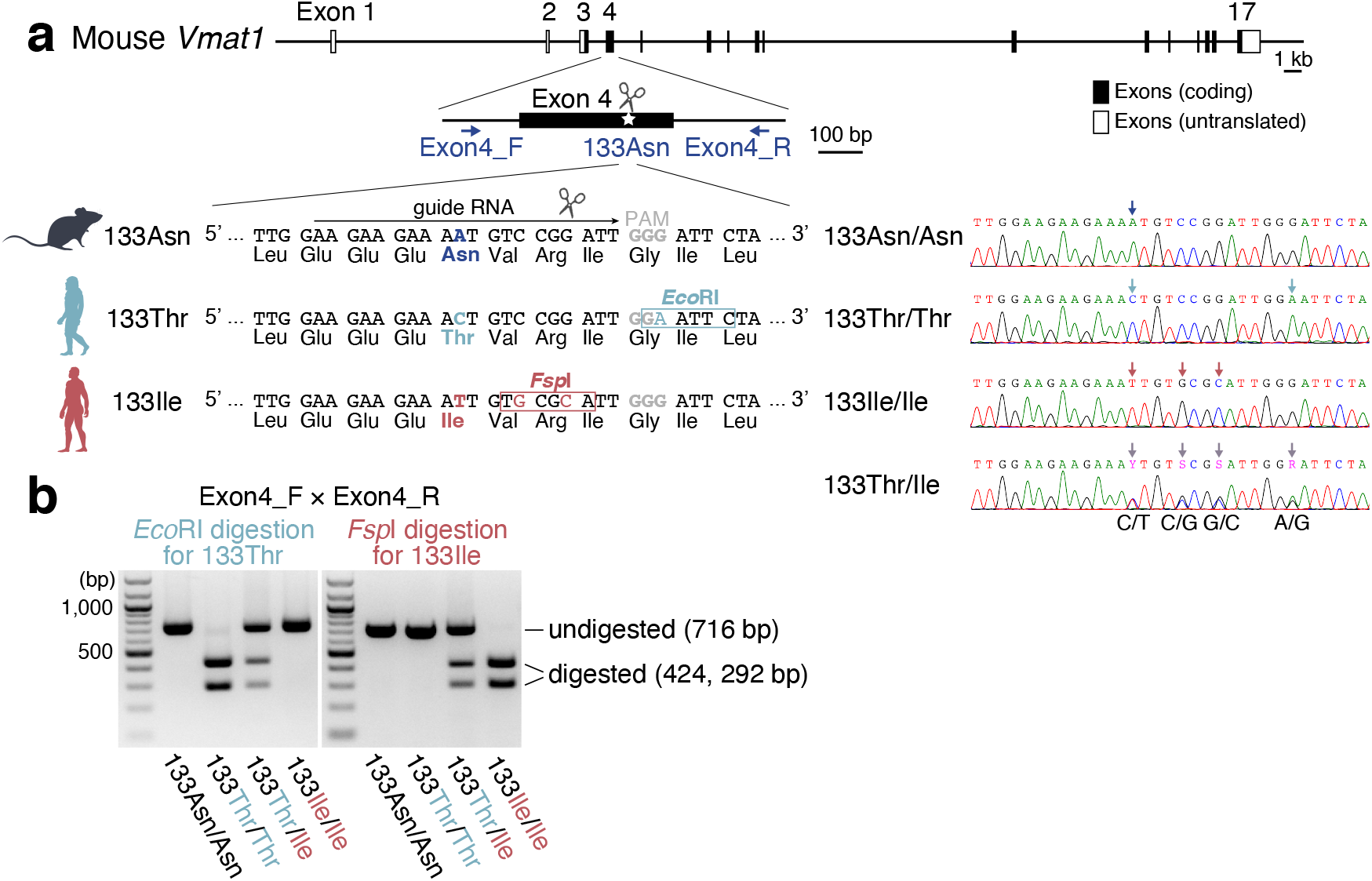
Generation of the *Vmat1*-humanized mouse models by CRISPR/Cas9 genome editing. (a) Targeting strategy for mVMAT1 133Asn humanization. Genetic configuration of the mouse *Vmat1* gene is shown above. Exon 4 encoding 133Asn is enlarged, and the primers used for genotyping (Exon4_F and Exon4_R) are depicted. To replace the mouse 133Asn with 133Thr or 133Ile by CRISPR/Cas9-mediated genome engineering, a guide RNA with minimum off-target effects was designed. GGG (gray) represents the PAM sequence. In addition to 133Asn humanization, restriction enzyme recognition sites (*Eco*RI and *Fsp*I) were synonymously incorporated to avoid the unwanted re-editing and to simplify genotyping. Sanger sequencing profiles of 133Asn/Asn (WT), 133Thr/Thr, 133Ile/Ile, and 133Thr/Ile are shown on the right. **(b)** PCR-RFLP assay, in which PCR products amplified using Exon4_F and Exon4_R were digested by *Eco*RI and *Fsp*I, respectively, could be used to distinguish 4 genotypes without sequencing.

### The human Ile substitution in mVMAT1 reduced behavioral anxiety

We accordingly conducted comprehensive behavioral tests among the 20 sets of mice including one of each genotype (*Vmat1*^WT^, *Vmat1*^Thr/Thr^, *Vmat1*^Thr/Ile^, and *Vmat1*^Ile/Ile^) from the 1st batch at 9–55 weeks (see Supplementary Table S3 for detailed information on age and the number of individuals used in each test). Detailed results were shown in Fig. S4–19. Tests of social interaction (SI) between stranger mice of the same genotype in a novel environment (one-chamber SI test) and of preference for a novel stranger over a previously exposed stranger (now familiar) in Crawley’s 3-chamber social interaction (CSI) test revealed marginal differences in behavioral phenotypes between mice with and without *Vmat1*^Ile^ (*P* = 0.062 for interactive effects of genotype (Ile) and place, generalized additive model (GAM) with quasi-Poisson distribution; Fig. 3a, Supplementary Fig. S9 and 10). In the CSI, *Vmat1*^Thr/Thr^ mice significantly preferred familiar to stranger mice (*P* = 3.26×10^-4^), while no such tendency was observed in *Vmat1*^Ile^ mice (*P* = 0.759 for *Vmat1*^Thr/Ile^ mice and (*P* = 0.195 for *Vmat1*^Ile/Ile^ mice; Fig. 3a). Moreover, we found consistently lower levels of anxiety among *Vmat1*^Ile^ mice across tests, including relatively greater preference for the (anxiogenic) light box during the two-chamber light/dark transition (LD) test and for the center of an open field (OF) (Supplementary Fig. S5 and 6).

**Figure 3.**
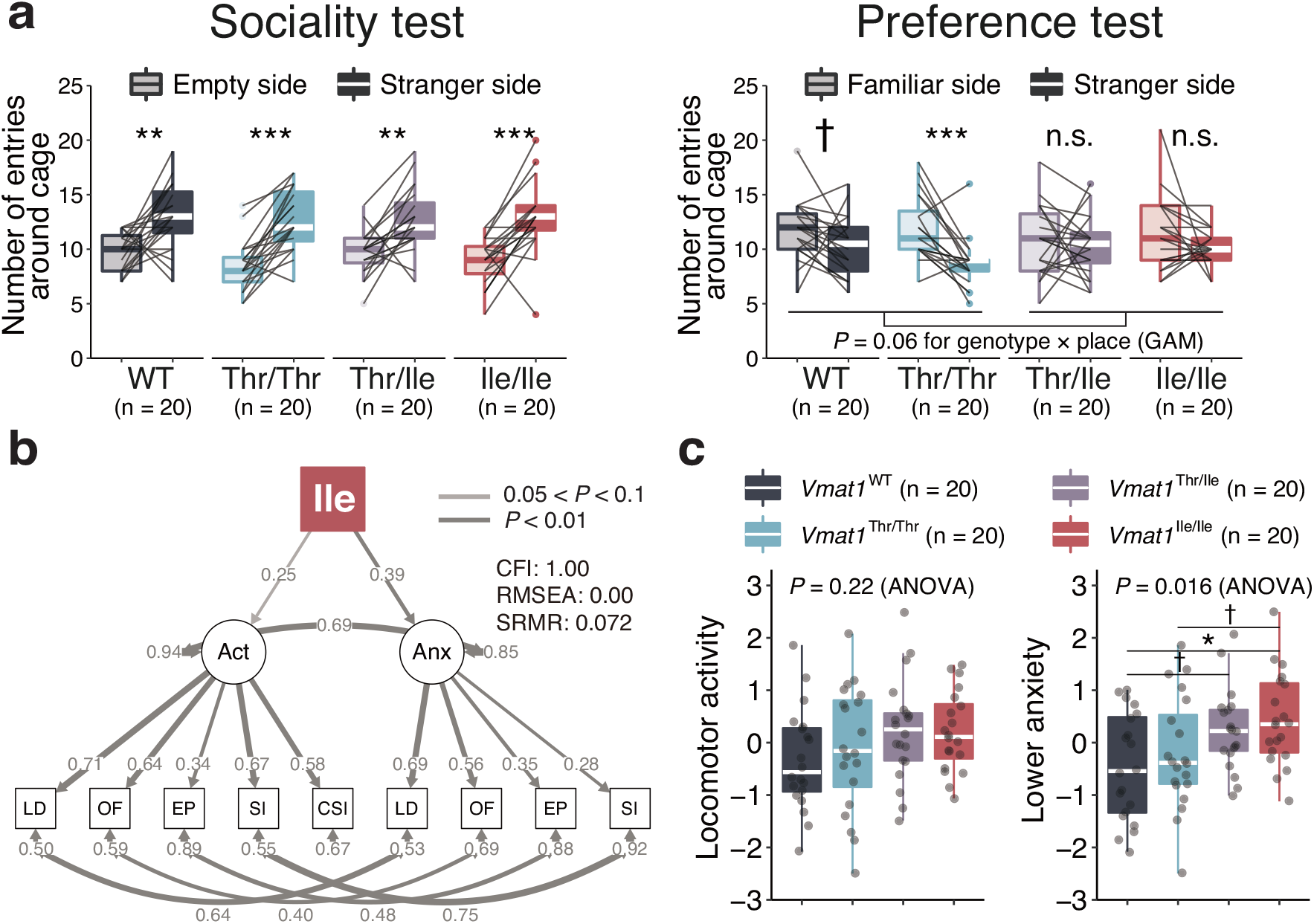
Comprehensive behavioral tests reveal distinct behavioral changes in *Vmat1*^Thr/Thr^ and *Vmat1*^Ile^ mice, including reduced anxiety. (a) In Crawley’s social interaction test, *Vmat1*^Thr/Thr^ mice preferred a familiar (previously exposed) mouse over a novel stranger. **(b)** Structural Equation Modeling (SEM) revealed the best fit model to explain the effects of *Vmat1* genotype on locomotor activity (Act) and anxiety (Anx). According to the model, *Vmat1*^Ile^ (*Vmat1*^Thr/Ile^ and *Vmat1*^Ile/Ile^) significantly reduces anxiety-like behavior. The boxes and circles represent measured and latent variables, respectively. The paths represent causal relationships on which numerical values indicate standardized coefficient and the gray-scale intensity of the paths indicates statistical significance tested by *t*-test. See Methods for the detailed modeling procedure. (c) *Vmat1*^Ile^ mice exhibited lower anxiety than WT and *Vmat1*^Thr/Thr^ genotypes. See Methods for the detailed calculation of the scores. For the composite anxiety score, a higher value indicates lower anxiety. Statistical significance was evaluated by paired *t*-test for **(a),** and by pair-wise *t*-test with FDR correction by the Benjamini-Hochberg method for **(c)**. Interactive effects of genotype (Ile allele labeled by 1, and 0 otherwise) and cage place were also assessed by generalized additive model with quasi-Poisson distribution in **(a)**. †: 0.05 < *P* < 0.1, *: 0.01 < *P* < 0.05, **: 0.001 < *P* < 0.01, ***: *P* < 0.001.

To confirm a pervasive effect on specific behavioral domains by humanized VMAT1, we constructed composite behavioral phenotype scores by summating standardized scores on related tests (LD, OF, elevated plus maze (EP), and SI tests) and compared the results among genotypes. We further conducted Structural Equation Modeling (SEM) to investigate the effects of genotype on these composite scores. This analysis revealed a significant effect of 133Ile on anxiety scores (*R*^2^ = 0.15, *P* = 8.7×10^-3^) but only marginal effects on locomotor activity (*R*^2^ = 0.062, *P* = 0.053; Fig. 3b).

Anxiety levels were generally lower in *Vmat1*^Ile^ mice (*P* = 0.016, one-way ANOVA; Fig 3c).

In addition to standard tests of anxiety, exploratory tendency, and locomotor activity, we also performed a novel delayed reward task to evaluate the differences in impulsivity between *Vmat1*^Thr/Thr^ and *Vmat1*^Ile/Ile^ mice (n = 10 for both genotypes, see Methods for detailed experimental procedure; Supplementary Fig. S19a–c). Throughout the test, there was no statistically significant difference in food preference between genotypes (*P* = 0.50 for the first term, GAM with quasi-Poisson distribution; Supplementary Fig. S19d). However, *Vmat1*^Ile/Ile^ showed a lower preference for the higher-calorie but delayed food reward than *Vmat1*^Thr/Thr^ mice every test day from 5 to 9 except for day 6 compared to the basal level after habituation (day 4) (Supplementary Fig. S19d).

### Differential regulation of dopaminergic and neurodevelopmental genes in the amygdala by *Vmat1* genotype

We then conducted RNA-Seq analysis to investigate the effects of the four *Vmat1* genotypes on gene expression patterns in three brain regions likely associated with the behavioral phenotypes (prefrontal cortex, amygdala, and striatum) (n = 4 mice for each genotype; see Supplementary Table S4 for detailed information on samples and reads). Principal component analysis revealed distinct expression patterns among brain areas, but no obvious differences between genotypes and ages (4 vs. 10 months) (Supplementary Fig. S20).

We thus screened for differentially expressed genes (DEGs) by pair-wise comparisons between genotypes, which yielded a total of 80 DEGs exclusively in the amygdala (Fig. 4a). A large proportion of these genes (56 out of 80) were differentially expressed between *Vmat1*^WT^ and *Vmat1*^Ile/Ile^ mice (Fig. 4a). The expression levels of DEGs for each sample were shown in Supplementary Fig. S21. We then utilized the correlations between individual composite anxiety scores and expression levels of the detected amygdalar DEGs among a subset of 8 mice (n = 2 for each genotype).

**Figure 4.**
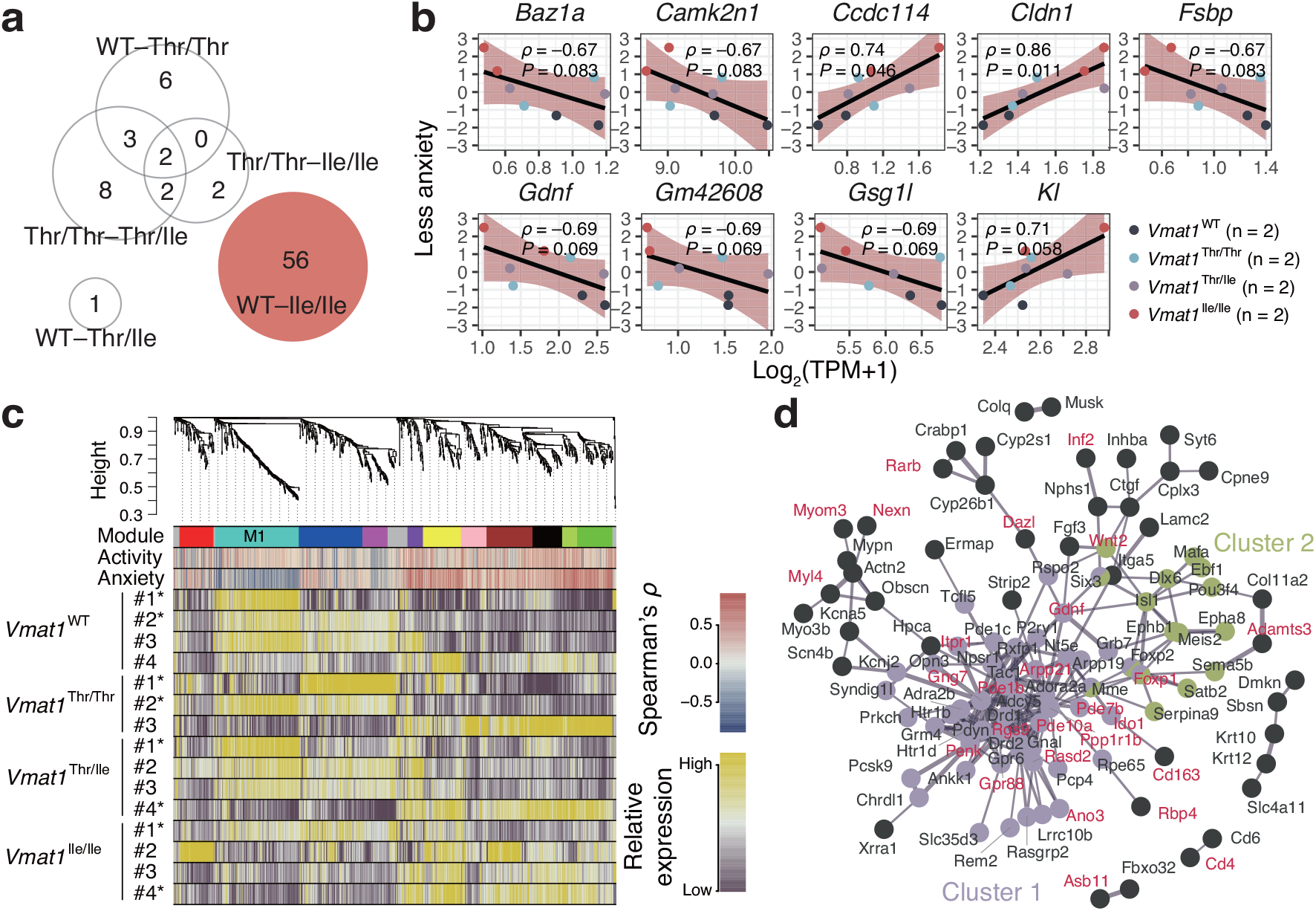
Differentially expressed genes (DEGs) in the brain among *Vmat1* genotypes and predicted co-expressing modules. **(a)** The number of DEGs detected by pair-wise comparisons among the four genotypes. All DEGs were found in the amygdala (with none in prefrontal cortex or striatum). **(b)** Correlations between individual DEG expression levels for the WT vs. and *Vmat1*^Ile/Ile^ comparison and composite anxiety scores from LD, EP, OF, and SI tests (see Methods). Only genes with strong Spearman’s correlations (*P* < 0.1) are shown. The bands are 95% confidence intervals. **(c)** Network dendrogram from co-expression modules based on the expression data of all 47 regional brain samples. Each branch represents an individual gene, and the colors below represent the module, correlation (ρ) with behavioral phenotype (locomotor activity and anxiety), and the relative expression level in the amygdala across genotypes. The samples with asterisks are from mice with behavioral data and were used to calculate the correlations between expression levels and behavioral phenotypes. The M1 module (shown in turquoise), showing negative correlations with anxiety score exhibited significant overrepresentation of the DEGs detected between WT and *Vmat1*^Ile/Ile^ mice. **(d)** Protein–protein interaction networks among the genes in M1. The DEGs detected between WT and *Vmat1*^Ile/Ile^ mice are shown in red. The thickness of the line indicates the strength of data support analyzed by STRING.

Particularly strong correlations were found for nine genes, *Baz1a*, *Camk2n1*, *Ccdc114*, *Cldn1*, *Fsbp*, *Gdnf*, *Gm42608*, *Gsg1l*, and *KI* (*P* < 0.1; Fig. 4b, Supplementary Fig. S22). The DEGs were further examined for gene ontology (GO) enrichment, which revealed significant overrepresentation of genes involved in “behavior”, “fear response”, “response to amphetamine”, “cardiac muscle tissue development”, or “cAMP catabolic process” (Table 2) between WT and Ile/Ile mice, while no GO terms were enriched in DEGs for other genotype comparisons (as there were few such genes). Moreover, gene expression patterns in the Ile mice amygdala were significantly positively correlated with those of striata of transgenic mouse models of Huntington disease (P < 1.0 × 10^-28^; Supplementary Table S5, Fig. S23).

**Table 2.**
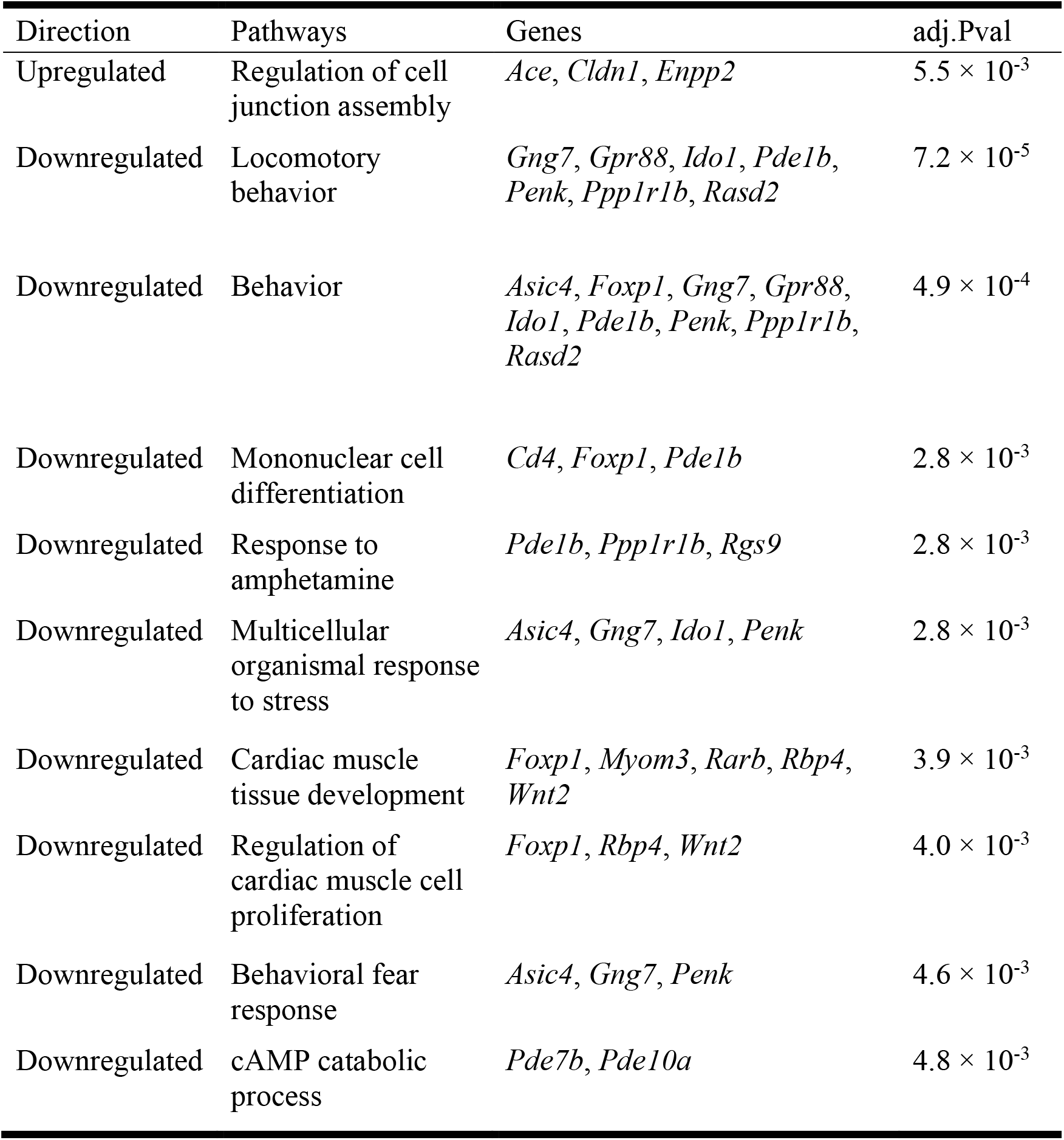
List of gene ontology (GO) terms significantly overrepresented in the set of differentially expressed genes (DEGs) between Vmat1^WT^ and Vmat1^Ile/Ile^ mice. P values were corrected by the Benjamini-Hochberg method. Only terms with *P* < 0.05 are presented. Downregulation indicates *Vmat1*^WT^ > *Vmat1*^Ile/Ile^ while upregulation indicates *Vmat1*^Ile/Ile^ > *Vmat1*^WT^.

To further characterize the biological pathways affected by *Vmat1* genotype, we performed weighted gene correlation network analysis (WGCNA; Fig. 4c). A co-expressing gene module (M1; Fig. 4c) stood out as it was significantly overrepresented with DEGs detected in the WT vs. Ile/Ile comparison (odds ratio = 3.79, *P* = 2.88×10^-8^ by Fisher’s exact test) and was negatively correlated with composite anxiety score.

These genes include many involved in adrenergic (*Adra2b*), dopaminergic (*Adcy5*, *Adora2a*, *Drd1*, *Drd2*, *Gnal*, *Gng7*, *Gpr6*, *Gpr88*, *Pde1b*, *Pde1c*, *Pde7b*, *Pde10a*, *Pdyn*, *Penk*, *Ppp1r1b*, *Rasgrp2*, *Rgs9* and *Tac1*), glutamatergic (*Grm4*), and serotonergic (*Htr1b*, *Htr1d*, *Htr1f*, and *Htr4*) signaling pathways. Sub-network analysis further detected another functional gene cluster involved in neural development within the co-expressing module (cluster 2 in Fig. 4d). Strikingly, three genes, *Foxp1*, *Foxp2*, or *Six3*, all of which are important regulators of neural growth, were located in hubs connecting genes involved in postsynaptic signaling (cluster 1) as mentioned above and neural development (cluster 2; e.g., *Dlx6*, *Ebf1*, *Isl1*, and Wnt2).

### The humanized mVMAT1 disrupted anxiety-associated theta band activity in the basolateral amygdala

Aforementioned behavioral and transcriptomic results suggest that the *Vmat1* genotype influences anxiety-related neuronal activity patterns. Inspired by a previous report that theta (4–12 Hz) oscillations in the medial prefrontal cortex (mPFC) and basolateral amygdala (BLA) are specifically enhanced under anxiogenic environments such as the open arms of an elevated plus maze (EP) and the center area of an OF (19, 20), we next performed simultaneous local field potential (LFP) recordings from the dorsomedial prefrontal cortex (dmPFC) and BLA in freely moving mice during elevated plus maze (EP) exploration (Fig. 5a; n = 11 dmPFC and 9 BLA electrodes from 5 *Vmat1*^WT^, n = 6 and 6 electrodes from 4 *Vmat1*^Thr/Thr^ and n = 10 and 7 electrodes from 4 *Vmat1*^Ile/Ile^ mice). The histological confirmation of electrode positions and raw LFP signals were shown in Fig. 5b and c, respectively. Power spectral analysis of WT mice revealed lower 4–7 Hz power in both dmPFC and BLA during exploration of the EP open arms compared to close arms, suggesting a relationship with anxiety-like behavior (Fig. 5d). Further, the Granger causality test revealed the directionality of 4–7 Hz oscillations from the mPFC to BLA (Fig. 5e; n = 20 electrode pairs from 5 *Vmat1*^WT^ mice), consistent with anatomical evidence that the dmPFC preferentially projects to the BLA (21, 22) and that mPFC theta oscillations drive BLA neuronal activity (20). Based on these observations, we compared 4–7 Hz power in dmPFC and BLA among genotypes. Both WT and *Vmat1*^Thr/Thr^ mice exhibited significantly lower LFP 4–7 Hz signal power in dmPFC and BLA under the anxiogenic condition of open arm exploration compared to closed arm exploration, while *Vmat1*^Ile/Ile^ mice showed no such consistent activity pattern in the BLA (Fig. 5f), suggesting that anxiety-related neuronal mechanisms in the dmPFC–BLA circuit are disrupted by the *Vmat1*^Ile/Ile^ genotype.

**Figure 5.**
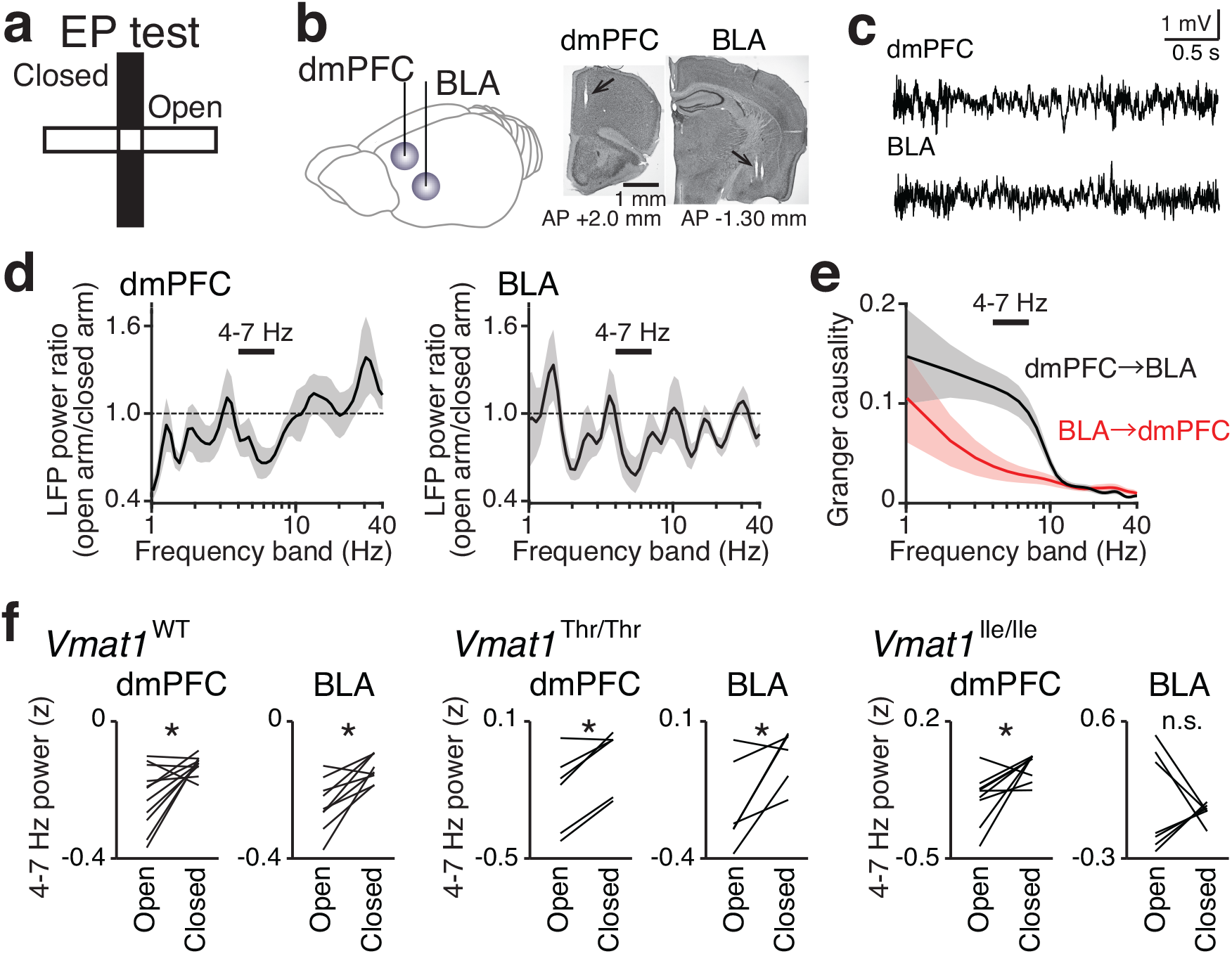
Wild-type and *Vmat1*^Thr/Thr^ mice, but not *Vmat1*^Ile/Ile^ mice, exhibit a reduction in amygdalar 4–7 Hz local field potential (LFP) power under anxiogenic conditions. (a) (Left) LFP recordings were simultaneously performed from the dorsomedial prefrontal cortex (dmPFC) and basolateral amygdala (BLA). (Right) Histological confirmation of electrode locations in the dmPFC and BLA. **(b)** Typical LFP signals from the dmPFC and BLA. **(c)** An elevated plus maze (EP) test. **(d)** Spectrograms of dmPFC (left) and BLA (right) LFP power in (anxiogenic) open arms relative to closed arms. Data were averaged from all *Vmat1*^WT^ mice. The bar above indicates the 4–7 Hz band, showing pronounced decreases in LFP power in both regions. **(e)** Spectral Granger causality averaged over 20 dmPFC–BLA electrode pairs. **(f)** Comparison of dmPFC and BLA LFP 4–7 Hz power (z-scored) between open and closed arms. \**P* < 0.05, paired *t*-test. Each line represents one electrode.

## Discussion

While the potential importance of *VMAT1* in neuroendocrine signaling has been suggested, few studies have focused on this gene until recently, largely due to its low expression in the central nervous system (12, 23, 24). A series of studies, however, have demonstrated the psychopathological effects of this gene and suggests its importance in emotional regulation. The Thr136Ile polymorphism (rs1390938) is allegedly associated with bipolar disorder (16), autism spectrum disorder (25), cognitive function related to schizophrenia (26), anxiety, depressiveness, neuroticism, or maladaptive impulsivity (18). Our previous study uncovered from an evolutionary perspective that this variant is a human-specific polymorphism (as shown in Fig. 1a) and under positive selection in the human lineage (11). Two substitutions (i.e., Glu130Gly and Asn136Thr) occurred in the human lineage after the divergence from the common ancestor of chimpanzees and humans, and these substitutions combined have been shown to decrease the monoamine uptake of VMAT1 (15). On the other hand, the new “hyper-function” allele (i.e. 136Ile; Lohoff et al. 2014), which increases the monoamine uptake of VMAT1 and is associated with fewer psychopathological symptoms by contrast, emerged just before the Out-of-Africa event of modern humans and is currently 20–61% in frequency in non-African populations (11). We also found that the Thr136Ile polymorphism has been under balancing selection in non-African populations (11). Given such evolutionary significance and psychopathological effects of the Thr136Ile polymorphism in *VMAT1*, this variant could play a very important role in exploring the evolutionary origins of human psychological traits and their diversity.

### Phenotypic changes in mice caused by the humanized *Vmat1*

Our comprehensive behavioral tests show the protective effects of the *Vmat1*^Ile^ allele against anxiety-like behavior (Fig. 3b and c), which is well consistent with the human phenotype (18). Compared to *Vmat1*^Ile^ (*Vmat1*^Thr/Ile^ and *Vmat1*^Ile/Ile^) mice, both wild-type and *Vmat1*^Thr/Thr^ mice showed similar behavioral phenotypes and have higher levels of anxiety in both solitary (LD, EP, and OF) and social (SI and CSI) environment. This is expected from the molecular evidence that 136Asn and 136Thr are comparable in the level of monoamine uptake, which is lower than that of 136Ile (15). It is noteworthy that the genotypic effect was not observed for fear response of mice (Supplementary Fig. S16). Given the potential difference in neurological mechanism behind fear and anxiety (27, 28), Vmat1 may specifically regulate anxiety-related circuits of the brain as discussed later. The results of delayed reward task may suggest that the *Vmat1*^Ile/Ile^ mice are more impulsive, at least initially, under this condition but contradicts an epidemiological study reporting an association of 136Thr with maladaptive impulsivity (18). Kim et al. (2018) reported that the deletion of the D2 dopamine receptor (D2R) increased impulsive behavior in mice, whereas restoration of D2R in the amygdala alone normalized impulsive behavior (29). This is compatible with our results, as dopaminergic genes including *Drd2* were downregulated in *Vmat1*^Ile/Ile^ mice (Supplementary Fig. S24). The possible difference in association of *VMAT1* genotype with impulsivity between humans and mice may stem from functional connectivity changes in related neural circuits and warrants further investigation. Overall, these observations suggest that the 136Ile substitution may contribute to uniquely human behavioral traits that have propelled both our technological progress and adaptation to almost all ecosystems, namely high exploratory tendency and relatively lower anxiety under threatening or novel conditions. Indeed, 136Ile allele frequency increases along the migration route of modern humans (11), suggesting that the exploratory tendency and/or boldness conferred by this mutation was advantageous for survival. In this regard, *Vmat1*^Ile^ is analogous to the D4R polymorphism associated with novelty-seeking behavior (30).

Differentially expressed genes were significantly enriched in those downregulated in *Vmat1*^Ile/Ile^ and involved in behavioral control and/or fear response as well Huntington disease (HD). It is possible that the Ile mice recapitulate some dimensions of HD pathology, given that dopamine (DA) signaling likely plays a significant role in HD, and the only current FDA-approved drug for HD is a VMAT2 inhibitor, tetrabenazine (31). WGCNA detected a co-expressing module (M1; Fig. 4c) in which the DEGs were significantly overrepresented and revealed that many genes involved in monoaminergic signaling pathways were co-expressed with the detected DEGs (Fig. 4d). Lohoff et al. reported that *Vmat1* KO affected DA signaling in particular, with upregulation of D2R and downregulation of tyrosine hydroxylase (TH) observed in both the frontal cortex and striatum (32). Given the association between *Vmat1* KO and decreased DA in the frontal cortex, we speculated that *Vmat1*^Ile/Ile^ mice would also exhibit downregulation of D2R and upregulation of TH compared to WT mice. Indeed, D2R (along with many other dopaminergic pathway genes) was downregulated in *Vmat1*^Ile/Ile^ mice compared to WT mice. However, there were no significant differences in TH expression among genotypes (Supplementary Fig. S24), suggesting that the *Vmat1* Ile allele alters dopaminergic transmission but not DA synthesis. The sub-network analysis detected another functional gene cluster involved in neural development within the co-expressing module (cluster 2 in Fig. 4d), such as *Foxp1*, *Foxp2*, *Six3*, *Dlx6*, *Ebf1*, *Isl1*, and *Wnt2*. Foxp1 (forkhead box protein P1) is a transcription factor that regulates the development of various tissues including the brain. Foxp1 heterodimerizes with Foxp2, and a human-specific Foxp2 substitution is associated with language impairment (33, 34). Foxp2 also regulates D2R expression (35) and extracellular DA levels (34), potentially involved in the neural development (36), while the cluster 2 genes *Dlx6*, *Ebf1*, *Isl1*, and *Six*3 are all implicated in the development of striatal neurons that express DA receptors (37–39). The DEG Wnt2 (also known as Irp) has been found to increase dopaminergic neurons by inducing the proliferation of progenitor cells in the developing midbrain (40). The differential expression of these genes may also be related to the regulatory role of VMAT1 in hippocampal neurogenesis (41), which emerging evidence suggests is associated with anxiety-like behavior (42, 43). Taken together, the differential regulation of DA signaling by *Vmat1* genotypes may have widespread effects on neural development and plasticity.

The electrophysiological analysis revealed abnormal neuronal activity in the amygdala of *Vmat1*^Ile/Ile^ mice under fearful conditions (i.e., open arms in EP test; Fig. 5f). This was not seen in the medial prefrontal cortex, and thus suggests that amygdala-specific neuronal disturbance underlies the reduced anxiety-like behavior associated with this genotype. This is consistent with a functional imaging study showing that 136Ile was associated with increased reactivity and decreased habituation of amygdala toward threat-related stimuli in humans (14). Additional studies are required to reveal the precise association between these LFP changes and amygdalar output in *Vmat1*^Ile^ mice. In light of our behavioral results, we suggest that *Vmat1*^Ile^ may promote the activity of neurons that dampen anxiety, such those projecting from the BLA to the central amygdala (44, 45).

### Amygdala-specific transcriptomic and neuronal changes associated with anxiety imply intra- and inter-species differences in emotion and social behavior

One of the major findings of this study is that *Vmat1* genotype affected transcriptomic regulation and neuronal activity primarily in the amygdala. Given that amygdalar DA signaling is a powerful modulator of fear and anxiety (46), the alterations in gene expression and oscillatory neuronal activity likely contribute to the observed reduction in anxiety-like behaviors among *Vmat1*^Ile^ mice compared to WT mice. The abnormal neuronal activity pattern detected in the amygdala but not PFC of *Vmat1*^Ile^ mice may stem from a difference in VMAT1 expression levels across the brain. Previous studies verified that *VMAT1* mRNA and protein expression levels are relatively high in the amygdala compared to other brain regions in both rats and humans (16, 47). Our transcriptome analysis also revealed higher expression of *Vmat1* in the amygdala than in the striatum (*P* = 0.0097, Dunnett’s test; Supplementary Fig. S25). In contrast, *Vmat2* expression was comparable across regions (Supplementary Fig. S25), indicating distinctive regional regulation between subtypes as shown in a previous study (41). However, the detailed transcriptomic effects of Vmat1 on various types of neurons within the amygdala and possible contributions of peripheral monoamine signaling to them are still unclear, and thus need to be investigated with greater spatial resolution in future studies.

Neuronal circuits within the amygdala store associations between environmental cues and adverse stimuli as changes in synaptic strength, and these modified circuits drive the expression of emotion-related behaviors and physiological responses. Thus, the amygdala is a critical regulator of emotional and social behaviors (48), and in fact amygdalar dysfunction is implicated in a number of neuropsychiatric disorders (49–51). Although it is an evolutionarily primitive brain region with relatively well conserved gross structure and general function across species (45), recent studies have provided evidence for more subtle structural and/or functional differences within and among species potentially linked to differential regulation of monoaminergic signaling via VMAT1. The size of the amygdala has been shown to correlate with creativity (52), mental state inference (53), and the size and complexity of social networks in humans (54) and non-human primates (55). The DA response in the medial amygdala network is associated with mother‒infant bonding (56). Collectively, these findings strongly suggest a substantial contribution of the amygdala to evolution of the human social brain by acting as a hub among brain networks associated with emotion, cognition, and communication (57).

### A single substitution in humanized mouse models highlight molecular and neurological evolution underlying human emotional traits

Genetic deletion (KO) and overexpression have become predominant strategies for examining the neurological, behavioral, and pathogenic functions of specific genes. However, gene deletion and overexpression may alter the expression levels of many additional genes, and thus the observed phenotype may be distinct from that induced by target-specific pharmacological manipulations. The present study employed an alternative “knock-in” strategy of a non-synonymous *Vmat1* mutation and succeeded in capturing moderate differences in transcriptomic, neuronal, and behavioral phenotypes. Noticeably, the variant evaluated is a human-specific substitution (i.e., Thr136Ile of *VMAT1*) under selection that is not possessed by other mammals (Fig. 1a). This knock-in model thus provides a unique opportunity to examine the molecular and neurological mechanisms that distinguish human behavior from that of other primates. Such extensive functional analysis of a single mutation has been limited to a few genes such as *FOXP2* (34, 58) and never for effects on emotional traits (59). Therefore, the present study is the first to suggest a new experimental strategy for studies on the evolution of human brain mechanisms underlying emotion and related behavioral traits.

Lastly, it needs to be noted that recent genome-wide association studies (GWAS) on psychiatric disorders such as depression (60) and schizophrenia (61), and on specific personality traits (62) have not detected VMAT1 as a top hit despite alleged associations with these phenotypes under study (18, 63). This may indicate that VMAT1 has a weaker influence on phenotype than other significant loci, or that this gene must interact with other loci (epistasis) and/or environmental factors (G × E interaction) for measurable effects on phenotype. Such dependence on other genes or external factors could obscure phenotypic associations of VMAT1 evaluated by conventional approaches of GWAS. In fact, we found that a single amino acid substitution (Asn133Ile) altered the expression of 56 genes within the amygdala, any or all of which may contribute to the observed changes in behavioral phenotype. The effects of various environmental factors (which were largely controlled in this experiment through group housing and shared dams) and DEGs on these behavioral and neurological phenotypes would warrant further study.

## Methods

Fully detailed and referenced methods are available in Supplementary Methods.

### Phylogenetic analysis and *in silico* prediction of mVMAT1 structure with humanized residues

A phylogenetic tree was constructed from 263 orthologous sequences of vertebrate *SLC18A1* (*VMAT1*), including an archaic hominin sequence that was constructed by replacing 136Ile in the human reference sequence with 136Thr. Homology modeling of the mouse VMAT1 (mVMAT1) protein structure was performed using the SWISS-MODEL (64) web server (http://swissmodel.expasy.org) with the visualization by using PyMOL 2.4.1 (DeLanoScientific, San Carlos, CA). Provean v1.1.3 (65) and SIFT (66) were used to estimate the intolerance for individual amino acid mutations introduced in mVMAT1 based on the evolutionary conservation and the chemical properties of the exchanged residues. DynaMut (67) was also used to evaluate the effects of humanized mutations on the stability and flexibility of VMAT1 protein structure.

### Generation of *Vmat1*-humanized mouse models by CRISPR/Cas9 genome editing

See Supplementary Methods for the detailed design and preparation of guide RNA and single-strand DNA (ssDNA) donors targeting mouse *Vmat1*. In addition to the intended humanization, the synonymous substitutions depicted in Fig. 2a were introduced near the 133rd site to prevent unwanted re-editing and enable PCR genotyping. The guide RNAs, Cas9 proteins, and the donor ssDNAs were electroporated into mouse zygotes following the standard protocol (68) to obtain founder knock-in mice (Supplementary Fig S1a). The 133Thr and 133Ile knock-in founders were screened by PCR-RFLP assay (Supplementary Fig. S1c), followed by Sanger sequencing to confirm the correct substitution (Supplementary Fig. S1d). To exclude the possible side effects from off-target cleavages, we sequenced 12 more potential off-target candidate loci predicted by CRISPOR (69) (http://crispor.org) (Supplementary Table S2, Supplementary Fig. S2).

After confirming the absence of off-target cleavages, 133Thr founder No.1 and 133Ile founder No.10 were crossed with WT C57BL/6J mice to obtain heterozygous F1 generations, which were again verified to possess the designed substitutions by Sanger sequencing. For behavioral tests, F5 or F6 homozygous mice were crossed to obtain Thr/Thr, Thr/Ile, and Ile/Ile genotypes. In addition, C57BL/6J WT males and females carrying the Asn/Asn genotype were crossed to supply control mice. To eliminate differences in rearing environment, newborn males of the 4 genotypes (Asn/Asn, Thr/Thr, Thr/Ile, and Ile/Ile) were grouped in sets and nursed by the same mothers.

### Animal care and experimental conditions for behavioral tests

Animals were housed in groups of four individuals of each genotype under a 12 h light/dark cycle (7:00 AM to 7:00 PM) with ad libitum access to food and water. Adult male mice were used in all tests to eliminate the behavioral effects of the estrus cycle. The composition of cohorts and ages of the mice for every experiment are summarized in Supplementary Table S3. All behavioral tests were conducted in a soundproof room, and as much effort as possible was spent on controlling the effects of confounding factors such as light intensity, temperature, and humidity for the tested group of cage mates. Furthermore, to minimize the impact of one test on subsequent tests in individual mice, behavioral assessments were conducted in the following order after general health and neurological screens: light/dark transition (LD), open field (OF), elevated plus maze (EP), hot plate, social interaction in a novel environment (SI), rotarod, Crawley’s 3- chamber social interaction test (CSI), startle response/prepulse inhibition, Porsolt forced swim, T-maze, Barnes maze, tail suspension, fear conditioning, and home cage social interaction tests. See Supplementary Methods for detailed descriptions of behavioral tests.

All animal care protocols and experiments were performed according to guidelines approved by the Animal Care and Use Committee of the National Institute of Neuroscience, National Center of Neurology and Psychiatry (approval numbers: 2017005, 2020007), the Animal Experiment Committee of Fujita Health University (approval number: APU19063), and the Experimental Animal Ethics Committees of Graduate School of Medicine, and the University of Tokyo (approval number: P29-14).

### Statistical analyses and Structural Equation Modeling (SEM)

In order to capture overall behavioral patterns among genotypes, we analyzed two major domains of mouse behavior, activity and anxiety, by standardizing, normalizing, and combining sets of related behavioral scores. Total distances traveled in LD, OF, EP, SI, and CSI were used as indices for activity, while time in the light compartment of the LD shuttle box, duration in the central area of the OF, time spent in open arms of the EP, and total duration of active contacts in the SI were used as indices of anxiety. To confirm the effects of *Vmat1* genotype on the behavioral composites and to enhance statistical power and improve reliability, we used Structural Equation Modeling (SEM) (70). The SEM model included a measurement model and a regression model. The measurement model consisted of two latent factors grouping tests of locomotor activity and anxiety-like behavior. The regression model evaluated the effect of genotype on the two latent factors. Goodness of model fit was evaluated by normed Comparative Fit Index (CFI), Root Mean Square Error of Approximation (RMSEA), and the Standardized Root Mean Square Residual (SRMR) and significance of paths was tested by *t*-test. Normalization of behavioral scores and SEM were performed using the *bestNormalize* and *lavaan* packages in R, respectively.

### *In vivo* electrophysiological recordings and histological analysis

Five male WT C57BL/6J mice (16–20 weeks old, SLC Shizuoka, Japan) with preoperative weights of 20–30 g as well as four male *Vmat1*^Ile/Ile^ mice (16–20 weeks old) and four male *Vmat1*^Thr/Thr^ mice (20–24 weeks old) obtained from the 2nd batch were implanted with intracranial electrodes for *in vivo* electrophysiological recordings. The animals were housed under a 12 h/12 h light/dark schedule with lights on at 7:00 AM prior to surgery. For local field potential (LFP) recordings, an array of 3 immobile tetrodes was stereotaxically implanted above the dmPFC (2.00 mm anterior and 0.50 mm lateral to bregma) at a depth of 1.40 mm and an array of 4 tetrodes was implanted in the amygdala (0.80 mm posterior and 3.00 mm lateral to bregma) at a depth of 4.40 mm using guide cannulae. Following surgery, each animal was housed in a separate transparent Plexiglas cage with free access to water and food for at least 7 days before recordings.

The mouse was connected to the recording equipment via a digitally programmable amplifier (Cereplex M, Blackrock Microsystems, Salt Lake City, UT) placed close to the animal’s head. Electrical signals during elevated plus maze (see methods for detail) were sampled at 2 kHz and low-pass filtered at 500 Hz with Cereplex Direct data acquisition system. The animal’s moment-to-moment position was tracked at 15 Hz using a video camera attached to the ceiling for 10 min. After a recording session, the mouse was overdosed with isoflurane, perfused intracardially with 4% paraformaldehyde (PFA) in phosphate-buffered saline (PBS, pH 7.4), and decapitated. The dissected brain was fixed overnight in 4% PFA/PBS and then cryoprotected by successive overnight incubations in 20% sucrose and 30% sucrose in PBS. Frozen coronal sections (100 μm) were cut using a microtome, mounted, and processed for cresyl violet staining. The positions of all electrodes were confirmed by identifying the corresponding electrode tracks in histological tissue sections.

### RNA-sequencing and transcriptome analysis

We obtained RNA-sequencing (RNA-seq) data from 3 tissues in the brain, prefrontal cortex, amygdala, and striatum of four mice per genotype, with DNBSEQ platform at the Beijing Genomics Institute (BGI, Hong Kong, China). The raw reads were quality checked, trimmed, and filtered using fastp 0.20.0 (71), yielding 20–21 million filtered reads per sample (Supplementary Table S4). STAR v2.7.5c (72) was then used for mapping the reads to the reference mouse genome (GRCm38.p6). We assigned reads to the Ensembl100 annotation and generated fragment counts by featureCounts (73), and the fragment counts were used to perform differential expression analysis with iDEP v0.91 workflow (74). After normalizing all counts according to effective library size, we retained genes with > 1 count per million (CPM) in half of the samples (24 samples). A total of 13,608 genes were then assessed in downstream analyses. Principal component analysis (PCA) was first performed to reveal the overall expression pattern of each sample. While most samples demonstrated distinct expression patterns, one supposed amygdala sample (derived from a 4-month-old *Vmat1*^Thr/Thr^ mouse) exhibited an expression pattern similar to the other striatal samples (Supplementary Fig. S20). Given the inconsistency in the expression pattern, we excluded this sample and conducted the normalization step again for the remaining 47 samples.

We conducted pair-wise between-genotype comparisons of the same brain regions to identify differentially expressed genes (DEGs), which were defined by |log2(fold change)| > 1 and false discovery rate (FDR; corrected by the Benjamini and Hochberg method) < 0.05. The DEGs were further characterized by enrichment analysis for gene ontology (GO) terms in the iDEP pipeline. An Illumina BaseSpace application (http://basespace.illumina.com/apps) was used to investigate the correlation in expression levels of DEGs among currently and previously collected datasets. To identify specific biological pathways affected by *Vmat1* genotypes, we conducted weighted gene correlation network analysis (WGCNA) (75) of 15 samples from the amygdala. The 1000 genes with most variable expression levels (largest coefficients of variation) among samples were retained, and the correlations among gene expression levels across samples were then calculated to detect co-expressing modules. We also calculated the correlations between the expression levels of a given gene in the amygdala and the behavioral composite (locomotor activity and anxiety), based on the data collected from 8 individuals (two for each genotype). Genes belonging to modules of interest were further included in protein-protein interaction networks constructed using STRING, and functional gene clusters were detected by the MCODE module of Cytoscape 3.8.2 (76).

## Supporting information

Supplemental methods, tables, and fitgues

## Acknowledgments

Some of the computations for transcriptomic analyses were performed on the NIG supercomputer at ROIS National Institute of Genetics. We thank Dr. Liu Jinsha and Ms. Eriko Koike for technical assistance with mouse embryo manipulations and care of newborns, Ms. Chikako Ozeki and Tamaki Murakami for their assistance with behavioral experiments, and Drs. Masayuki Koganezawa and Noriko Osumi for their valuable comments on the manuscript. This work was supported by the Japan Society for the Promotion of Science (Grants-in-Aid for Scientific Research 17H05934, 19H04892, and 16H06276 (AdAMS) to MK, 20J12055 to DXS, 17H05939 and 19H04897 to TS, 17H05967 and 19H04922 to YUI, and 16H06276 to TM) and Intramural Research Grants for Neurological and Psychiatric Disorder of NCNP (27-7, 30-9, and 3-9 to TI). This work was also supported by MEXT Promotion of Distinctive Joint Research Center Program (Grant Number JPMXP0618217663).

## Author contributions

DXS and MK conceived and DXS, YUI, SH, TK, and TS designed the study. DXS, YUI, TI, TM, TS, and MK acquired funding. YUI and YM conducted genome-editing experiments with mice. DXS, SH, KN, and TK designed and DXS and SH performed behavioral tests. DXS analyzed behavioral test results with the help of SH and GS. NK and TS performed electrophysiological and histological experiments. TS dissected all the brain samples for RNA-Seq, for which DXS and HH performed data analysis. DXS, YUI, TI, SH, TS, and MK wrote the manuscript with contributions and revisions from all authors. All authors have read and approved the final manuscript.

