## Supplemental methods, tables, and fitgues for "Humanized substitutions of *Vmat1* in mice alter amygdala-dependent behaviors associated with the evolution of anxiety"

This file includes:

- Supplementary Methods
- Tables S1 to S5
- Figures S1 to S25
- References

### Supplementary Methods

#### Multiple sequence alignments and phylogenetic tree of *VMAT1*

Two hundred sixty-three orthologous sequences of vertebrate *SLC18A1* (*VMAT1*) were obtained from Ensembl101. After aligning the sequences using webPRANK (1) with default parameters, we calculated the proportions of amino acids identical to the human *VMAT1* within aligned protein regions. Sequences with < 50% identity were then excluded. In the case of multiple sequences for a given species, that with the highest proportion of identical amino acids to human *VMAT1* was . A phylogenetic tree was then constructed from the 236 species sequences with > 50% identity to human *VMAT1* using IQTREE v2.1.1 (2) with the "-nt AUTO" setting, including the Neanderthal sequence, which we constructed by replacing 136Ile in the human reference sequence with 136Thr. Phylogenetic relationship was visualized using iTOL v5.6.3 (3) through a web interface (https://itol.embl.de). Animal silhouettes were obtained from phylopic (http://phylopic.org/) under Public Domain license.

#### *In silico* prediction of mVMAT1 structure and tolerance of mutated residues

Homology modeling of the mouse VMAT1 (mVMAT1) protein structure was performed using the SWISS-MODEL (4) web server (http://swissmodel.expasy.org). The human organic anion transporter MFSD10 (PDB: 6s4m.1.A) was selected as the best template, with sequence identity of 17.04 (verified June 2020) and QMEAN score of –8.40 according to SWISS-MODEL. The predicted 3D structure of mVMAT1 was visualized using PyMOL 2.4.1 (DeLanoScientific, San Carlos, CA). Provean v1.1.3 (5) and SIFT (6) were used to estimate the intolerance for individual amino acid mutations introduced in mVMAT1 based on the evolutionary conservation and the chemical properties of the exchanged residues. The computations were made using the Provean web server (http://provean.jcvi.org/protein_batch_submit.php?species=mouse; last accessed June 2020). A mutation is predicted as deleterious (i.e., likely to affect its protein function) when the Provean score is smaller than −2.5 or the SIFT score is smaller than 0.05. Furthermore, we used DynaMut (7) to evaluate the effects of humanized mutations on the stability and flexibility of VMAT1 protein structure. Briefly, DynaMut enables accurate assessment of mutation impact on protein stability by implementing and integrating well established normal mode approaches with graph-based signatures in a consensus predictor (7). Here we focused on two measurements: 1) folding free energy (ΔΔG), an index of the difference in stability between WT and mutant proteins in which positive values represent increased stability of the mutant, and 2) Δ vibrational entropy energy (ΔΔSVib ENCoM), a per-site index of the difference in flexibility between WT and mutant proteins. Calculations were conducted using a web server (http://biosig.unimelb.edu.au/dynamut/).

**Generation of *Vmat1*-humanized mouse models by CRISPR/Cas9 genome editing**

*Preparation of the CRISPR components and donor DNAs*

The target region within the mouse *Vmat1* exon 4 was first analyzed by using the web-based CRISPR design tool CRISPOR (8) (http://crispor.org), to select a guide RNA sequence for replacing mouse 133Asn with 133Thr or 133Ile via the CRISPR/Cas9 system with minimal risk of off-target cleavages. Then, two parts of the CRISPR guide RNA, crRNA and tracrRNA (shown in Fig. 2a, Supplementary Fig. S1a, and Supplementary Table S1), were chemically synthesized and purified by reversed-phase column chromatography (FASMAC, Japan). Recombinant Cas9 protein (EnGen Cas9 NLS from *Streptococcus pyogenes*) was purchased from New England Biolabs (Ipswich, MA). The single-strand DNA (ssDNA) donors containing the humanized substitution and the restriction enzyme recognition sites flanked by 42–50 bp homology arms (Supplementary Table S1) were subsequently designed and chemically synthesized by Eurofins Genomics (Tokyo, Japan). In addition to the intended humanization, the synonymous substitutions depicted in Fig. 2a were introduced near the 133rd site to prevent unwanted re-editing. For 133Thr substitution, an *Eco*RI site was synonymously integrated into the protospacer adjacent motif (PAM) sequence to be destroyed. An *Fsp*I site was similarly incorporated within the guide RNA target sequence for the 133Ile allele. These synonymous substitutions also made the genotyping procedure much easier by eliminating the need for laborious sequencing (Fig. 2b).

*Electroporation of mouse one-cell embryos*

One-cell-stage embryos were collected by *in vitro* fertilization using C57BL/6J super-ovulated females (Charles River, Japan) and stud males. The guide RNAs, Cas9 proteins, and the donor ssDNAs were electroporated into mouse zygotes following the standard protocol (9) to obtain founder knock-in mice (Supplementary Fig S1a).

*PCR Genotyping and sequencing analyses*

Genomic DNAs were prepared from the newborn mouse tail by treatment with proteinase K in Lysis Buffer. The 133Thr and 133Ile knock-in founders were screened by PCR-RFLP assay (Supplementary Fig. S1c). Briefly, PCR products were amplified using the primer pairs depicted in Fig. 2a (also listed in Supplementary Table S1) and then digested by *Eco*RI or *Fsp*I, respectively. For founders and F1 generations, PCR products were also analyzed by Sanger sequencing to confirm the correct substitution allele. After that, PCR-RFLP could be used reliably for genotyping of subsequent generations.

*Off-target analyses of founder mice*

To exclude the possible side effects from off-target cleavages, the genomic DNAs extracted from the founder mice (133Thr No.1 and 133Ile No.10, shown in Supplementary Fig. S1c and d) were analyzed by Sanger sequencing. We employed CRISPOR (8) (http://crispor.org) to predict potential off-target candidate loci, and the 12 loci identified (Supplementary Table S2) were amplified by PCR using the primers listed in Supplementary Table S1 and sequenced (results are shown in Supplementary Fig. S2).

*Preparation of humanized mouse models for behavioral tests, transcriptome analyses, and neurophysiological analyses*

After confirming the absence of off-target cleavages, 133Thr founder No.1 and 133Ile founder No.10 were crossed with WT C57BL/6J mice to obtain heterozygous F1 generations, which were again verified to possess the designed substitutions by Sanger sequencing. Homozygous 133Thr and 133Ile mice after F2 generations were then selected and maintained. For behavioral tests, F5 or F6 homozygous mice were crossed to obtain Thr/Thr, Thr/Ile, and Ile/Ile genotypes. In addition, C57BL/6J WT males and females carrying the Asn/Asn genotype were crossed to supply control mice. To eliminate differences in rearing environment, newborn males of the 4 genotypes (Asn/Asn, Thr/Thr, Thr/Ile, and Ile/Ile) were grouped in sets and nursed by the same mothers. At 4 weeks old, 20 sets of 4 males, one of each genotype, were weaned and housed in 20 separate cages. Behavioral tests were conducted between 9 and 55 weeks of age as indicated (Supplementary Table S3). A subset of these mice (1st batch) was also used for transcriptome analyses at 42–45 weeks. Likewise, 8 sets of mice (2nd batch) including all 4 genotypes were generated and used for transcriptome analysis at 15 weeks and electrophysiological recordings at 16–24 weeks.

**Behavioral battery tests**

*General health and neurological screens (GHNS)*

The presence of whiskers and bald patches was checked daily to ensure the health status of mice (10). The righting, whisker touch and ear twitch reflexes were also evaluated to insure age-appropriate neurological function. Body weight and rectal temperature were measured. In addition, neuromuscular function was assessed by grip strength and wire hanging tests. Briefly, grip strength was measured using a grip strength meter (O'Hara & Co., Tokyo, Japan). The mouse was positioned to spontaneously grasp a wire grid by the forelimbs and then pulled backward by the tail until wire release. The peak force was recorded in Newtons (N). Each mouse was tested three times, and the greatest value obtained was used for further analyses. In the wire hanging test, a mouse was placed on a wire mesh at the top of the apparatus (O'Hara & Co.), and the wire mesh was then gently turned upside down. The mouse gripped the wire in order not to fall off, and the latency to fall was recorded with a 60 s cut-off time.

*Light/dark transition (LD)*

The light/dark transition test was conducted to measure anxiety level as previously described (11). The apparatus consisted of two equal-sized plastic boxes (20 × 20 cm), one illuminated (390 ± 20 lx) and the other dark (< 2 lx), separated by a central partition plate with a small 3 × 5 cm opening allowing the mouse to transit from one box to the other. A mouse was first placed in the dark box and allowed to freely explore the apparatus for 10 min. The time to first entry into the light box, time spent in the light box, number of transitions between boxes, and distances traveled in light and dark boxes were measured using Image LD software (see "Image analysis of behavioral tests").

*Open field (OF)*

Locomotor activity and explorative tendency were measured using an open field apparatus (40 × 40 × 30 cm; Accuscan Instruments, Columbus, OH) as described previously (12). The test chamber was illuminated at 100 ± 5 lx. A mouse was placed in the corner of the apparatus and total distance traveled (cm), time spent in the center area (20 × 20 cm), vertical activity, and stereotypic counts over 120 min were recorded by the VersaMax system and analyzed using Image OF software (see "Image analysis of behavioral tests").

*Elevated plus maze (EP)*

An elevated plus-maze test was conducted as previously described (13). The apparatus consisted of two open arms (25 cm long × 5 cm wide) crossing two enclosed arms of identical dimensions but with 15-cm high transparent walls, all connected by a central platform. The entire apparatus was elevated 50 cm above the floor and illuminated uniformly at 100 lx. A mouse was placed at the center of the maze, facing one of the closed arms and allowed to freely explore for 10 min. Number of open and closed arm entries, distances traveled, proportions of entries into open arms, and proportions of time spent in open arms were measured using Image EP software (see "Image analysis of behavioral tests").

*Hot plate (HP)*

Pain sensitivity was measured by the hot plate test. A mouse was placed on a 55°C hot plate (Columbus Instruments, Columbus, OH), and latency to escape from the plate was recorded with a 15-s cut-off time.

*Social interaction in a novel environment (SI)*

The social interaction test in a novel environment test (one-chamber social interaction test) was performed as described previously (10). Two mice of the same genotype but reared in different cages were placed together in a box (40 × 40 × 30 cm) and allowed to explore freely for 10 min. Mouse behaviors were recorded with a CCD camera. The total duration of contact, total number of contacts, total duration of active contact, mean duration per contact, and total distance traveled were measured automatically using Image SI software (see "Image analysis of behavioral tests"). Active contact was defined as maintained contact while either mouse traveled more than 10 cm between two successive image frames acquired once per second.

*Rotarod (RR)*

Motor coordination and motor learning were examined using an accelerating rotarod (UGO Basile, Comerio, VA, Italy). The latency to fall from the rod was recorded during three daily trials conducted on 2 consecutive days (3 × 2 trials). During each trial, rod speed was increased from 4 to 40 rpm over the 5-min test period.

*Crawley's 3-chamber social interaction (CSI)*

Crawley's 3-chamber social interaction test was performed to measure sociability and social novelty preference (12). The experimental apparatus was a 41 × 62 cm rectangular non-transparent gray Plexiglas box separated into three equal-sized chambers (20 × 40 cm) by transparent Plexiglas plates with small openings allowing mice to freely transit from one chamber to another. One wire cylinder-shaped cage was placed in the corner of each side chamber. Tested mice (aged 16–19 weeks, see Table S2) were first allowed to freely explore the chambers for 10 min as a habituation period. A stranger mouse was then placed randomly in one of the side-chamber wire cages and the tested mouse was again allowed to freely explore the chambers for 10 min. In a final step, a second stranger mouse was placed in the wire cage in the opposite side chamber, and the tested mouse was again allowed to freely explore the chambers for 10 min. Number of contacts and interaction times with strangers 1 and 2 were recorded using Image CSI software (see "Image analysis of behavioral tests").

*Startle response / prepulse inhibition (PPI)*

Sensory-motor integration was tested by a prepulse inhibition test (10), which measures the effect of an audible pre-tone (prepulse) of variable intensity on the startle response to a louder pulse. The system used for the detection of startle reflexes (O'Hara & Co.) consisted of four standard cages placed on a movement sensor within sound-attenuated chambers with fan ventilation. Before each testing session, acoustic stimuli and mechanical responses were calibrated using devices supplied by the manufacturer. A test session was composed of 6 different trial types: two startle stimulus-only trials (110 or 120 db) and four prepulse inhibition trials (74 or 78 db prepulses delivered prior to a 110 or 120 db startle). Six blocks of the six trial types (i.e., 36 trials in total) were conducted, with each trial type presented once in pseudorandom order within each block. The PPI (%) was calculated for each trial type according to the following equation: {[(startle amplitude of trial without prepulse) – (startle amplitude of trial with prepulse)]/(startle amplitude of trial without prepulse)} × 100.

*Porsolt forced swim (PS)*

The Porsolt swim test was performed as previously described (10). The apparatus consisted of four Plexiglas cylinders (12 cm diameter × 22 cm height) filled with water (room temperature) up to a height of 7.5 cm. Four mice (a set) were placed individually in each cylinder, and images were captured at two frames per second for 10 min. Swimming distance and immobility time (% of total) were recorded automatically using Image TS software (see "Image analysis of behavioral tests").

*Tail suspension (TS)*

Mice were suspended 30 cm above the floor in a visually isolated area by adhesive tape placed approximately 1 cm from the base of tail. Behavior during suspension was recorded for 10 min using Image TS software (see "Image analysis of behavioral tests").

*T-maze (TM) forced alteration*

A T-maze forced alternation task was performed as previously described (14). The T maze was constructed of three white plastic runways with walls 25-cm high (O'Hara & Co.) partitioned into 6 areas by sliding doors. The stem of the "T" was divided into a start compartment area S1 at the intersection with the arms and a main area S2 (13 × 24 cm). Similarly, the T arms were divided into passageway areas P1 and P2 with S1 and main areas A1 and A2 (11 × 20.5 cm). Each trial consisted of one forced run followed by one free run. First, the mouse was placed in S2 and forced to proceed in one direction (S2 → A2 → P2 → S1, or S2 → A1 → P1 → S1) by opening the doors successively. After returning to S1 and a set inter-trial delay, mice were then allowed to choose to enter A1 or A2. If the mouse chose the opposite direction from the previous forced run, it was counted as a correct response, while if the mouse chose the same direction as the previous run, it was counted as an incorrect response. After 30 practice trials (10 per day for 3 consecutive days), trials with delay (4 trials with 3-s delays and 2 trials each with 30-, 60-, and 120-s delays) were conducted for 3 consecutive days. Data were acquired and sliding doors controlled automatically using Image TM software (see "Image analysis of behavioral tests").

*Barnes maze (BM)*

Spatial learning, spatial memory, and behavioral flexibility were tested using the Barnes maze as described previously (12). The maze was a circular whiteboard (1m in diameter) with 12 holes equally spaced around the margin and elevated 81.5 cm from the floor (O'Hara & Co.). A black plastic box (17 × 13 × 7 cm) lined with paper cage bedding was positioned under one of the holes (the target). The board position of the target (1–12) was constant for a given mouse but randomly assigned across individuals. The apparatus was illuminated at 800 lx or more. The board was rotated daily so that the spatial location of the target changed relative to distal visual room cues while proximal cues were held constant. A training trial was conducted per day over 20 days until 24 trials were completed in total before the 1st probe test. Each trial lasted a maximum of 5 min, and was completed when the mouse entered into the black plastic box. Probe tests were conducted for 3 min without the black plastic box one day or one month after the final training trial. The number of errors, latency and distance traveled to reach the target hole for the first time, the number of omission errors and time spent around each hole were recorded by Image BM software (see "Image analysis of behavioral tests").

*Fear conditioning (FZ)*

Contextual and cued fear conditioning tests were conducted and analyzed as described in a previous study (10). Mice were placed in a transparent acrylic chamber (26 × 34 × 33 cm, O'Hara & Co.) with 55 dB ambient white noise and 100 lx illumination. In the conditioning session, the conditioned acoustic stimulus (CS, 55 dB) was presented for 30 s at three times (2, 4, and 6 min), and a mild foot shock (2 s, 0.3 mA) was presented as the unconditional stimulus (US) at the end of each CS. One day after the conditioning session, contextual fear was measured in the same chamber, while cued fear was measured thereafter in a triangular box (33 × 33 × 33 cm) constructed of white opaque Plexiglas. Mice were placed in the chamber for 3 min with neither CS nor US presented.

Thereafter, the CS (55 dB) was presented for the last 3 min and images were captured at 1 frame per second. Each pair of successive frames in which the mouse moved was measured. When this measure was below a certain threshold (20 or 30 pixels), the mouse was considered to be "freezing"; alternatively, when this measure equaled or exceeded the threshold, the behavior was considered "non-freezing". "Freezing" that lasted less than 2 s was not included in the analysis. Data acquisition, control of stimuli (i.e., tones and shocks), and data analysis were conducted automatically using Image FZ software (see "Image analysis of behavioral tests").

*Home cage social interaction test (HCSI)*

The 24-hour home cage locomotor activity and social interaction test was conducted as previously described (15) for one continuous week. The monitoring system included a standard home cage (29 × 19 × 13 cm) with filtered cage top and an infrared video camera (O'Hara & Co.). Two mice of the same genotype that had been housed separately were placed together in the home cage and locomotor activity and social behavior monitored by video. Social interaction was measured by counting the number of "particles" (tracers for each mouse) detected in each frame, with two separate particles indicating no contact and one particle indicating physical contact (i.e., because the tracking software could not distinguish two separate bodies). Data acquisition and analysis were conducted using Image HA software (see "Image analysis of behavioral tests")

**Delayed reward task**

Following these comprehensive behavioral tests, we conducted an original delayed reward task to evaluate the impulsivity of *Vmat1*$^{Thr/Thr}$ and *Vmat1*$^{Ile/Ile}$ mice (n = 10 for both genotypes). In brief, a mouse was housed in a cage (22.5 × 32.5 × 21 cm; FDB-300W/FDL-8D, Melquest, Toyama, Japan) with controlled access to two separate feeding trays for 16 days. Consumption of normal food (3.57 kcal/g; CRF-1, Oriental Yeast Co., Ltd., Tokyo, Japan) and high-calorie food (Banana chip containing 5.42 kcal/g; 4515996091582, KALDI COFFEE FARM, Tokyo, Japan), considered small and large rewards, respectively, were measured daily by weighing the feeding trays. After habituating mice to this system for 4 days with ad libitum access to both foods, we programmed feeding tray access so that the normal food was provided for the first 45 min and high-calorie food provided only for the last 15 min of the three 1-h daily feeding time (7:00–8:00 PM, 10:00–11:00 PM, and 1:00–2:00 AM) for 6 consecutive days. Under such delayed reward conditions, consumption of the lower-calorie food is considered an impulsive behavior (i.e., does not maximize reward). Furthermore, to maximize the impulsivity, we set a fasting day after the 6 days of delayed feeding while monitoring animal weights (so as not to jeopardize the animal's health) and conducted the same test again for 5 days.

**Image analysis of behavioral tests**

We developed in-house programs to analyze the imaging data acquired for many of these behavioral tests (Image LD, Image OF, Image EP, Image SI, Image CSI, Image

TS, Image TM, Image BM, Image FZ, and Image HA) based on ImageJ (U.S. National Institutes of Health; available at https://imagej.nih.gov/ij/) and modified for each test by Tsuyoshi Miyakawa (available through O'Hara & Co.).

**Statistical analyses and Structural Equation Modeling (SEM)**

Group means were compared by paired sample *t*-test, unpaired Student's *t*-test, one-way ANOVA, or pair-wise *t*-test with FDR correction by the Benjamini-Hochberg method as indicated using R 4.0.2. We obtained comprehensive metrics for two major domains of mouse behavior, activity and anxiety, by standardizing, normalizing, and combining sets of related behavioral scores. Total distances traveled in LD, OF, EP, SI, and CSI were used as indices for activity, while time in the light compartment of the LD shuttle box, duration in the central area of the OF, time spent in open arms of the EP, and total duration of active contacts in the SI were used as indices of anxiety. To further confirm the effects of *Vmat1* genotype on the behavioral composites and to enhance statistical power and improve reliability, we used Structural Equation Modeling (SEM) (16). The SEM model included a measurement model and a regression model. The measurement model consisted of two latent factors grouping tests of locomotor activity and anxiety-like behavior, which were assessed by the same metrics above. The regression model evaluated the effect of genotype on the two latent factors. In the best fitted model, genotype was labeled by the presence of Ile allele as follows: $Vmat1^{WT}$ = 0, $Vmat1^{Thr/Thr}$ = 0, $Vmat1^{Thr/Ile}$ = 1, and $Vmat1^{Ile/Ile}$ = 1. Goodness of model fit was evaluated by normed Comparative Fit Index (CFI), Root Mean Square Error of Approximation

(RMSEA), and the Standardized Root Mean Square Residual (SRMR) and significance of paths was tested by *t*-test. Normalization of behavioral scores and SEM were performed using the *bestNormalize* and *lavaan* packages in R, respectively. Availability of the data and codes used are described in the section **Availability of data and materials**.

***In vivo* electrophysiological recordings**

*Subjects*

Five male WT C57BL/6J mice (16–20 weeks old, SLC Shizuoka, Japan) with preoperative weights of 20–30 g as well as four male *Vmat1*$^{Ile/Ile}$ mice (16–20 weeks old) and four male *Vmat1*$^{Thr/Thr}$ mice (20–24 weeks old) obtained from the 2nd batch were implanted with intracranial electrodes for *in vivo* electrophysiological recordings. The animals were housed under a 12 h/12 h light/dark schedule with lights on at 7:00 AM prior to surgery.

*Surgery*

Animals were anesthetized with isoflurane gas (1%–3%) and 1-mm diameter circular craniotomies were made using a high-speed drill at the target coordinates. For local field potential (LFP) recordings, an array of 3 immobile tetrodes was stereotaxically implanted above the dmPFC (2.00 mm anterior and 0.50 mm lateral to bregma) at a depth of 1.40 mm and an array of 4 tetrodes was implanted in the amygdala (0.80 mm posterior and 3.00 mm lateral to bregma) at a depth of 4.40 mm using guide cannulae.

The tetrodes were constructed from 17-μm diameter polyimide-coated platinum–iridium (90/10%) wire (California Fine Wire, Grover Beach, CA), and the individual electrode tips were plated with platinum to lower the impedance to 200–250 kΩ. Stainless steel screws were implanted on the skull attached to the brain surface to serve as ground/reference electrodes. To prevent brain tissue drying at the implant sites, a dummy cannula was inserted through the guide cannula and both covered by a cap. The entire recording device was secured to the skull using stainless steel screws and dental cement. After all surgical procedures were completed, anesthesia was discontinued, and the animals were allowed to awaken spontaneously. Following surgery, each animal was housed in a separate transparent Plexiglas cage with free access to water and food for at least 7 days before recordings.

*Electrophysiological recording*

The mouse was connected to the recording equipment via a digitally programmable amplifier (Cereplex M, Blackrock Microsystems, Salt Lake City, UT) placed close to the animal's head. For recording electrophysiological signals, the EIB of the microdrive array was connected to a Cereplex M digital headstage, and the digitized signals were transferred to a Cereplex Direct data acquisition system. Electrical signals were sampled at 2 kHz and low-pass filtered at 500 Hz. The animal's moment-to-moment position was tracked at 15 Hz using a video camera attached to the ceiling.

*Elevated plus maze (EP)*

The elevated plus maze used during electrophysiological recording was made of ABS resin and consisted of a central square (7.6 × 7.6 cm) and four arms (each 28 cm long × 7.6 cm wide), two open arms with no railing and two closed arms enclosed by transverse walls (15 cm in height). The maze was elevated 30 cm from the floor and illuminated with two 32-W fluorescent overhead lights, which produced light intensities of 250 ± 20 lx and 170 ± 20 lx in the open and closed arms, respectively. In a recording session, a mouse was placed in the center of the central square facing an open arm and allowed to explore the maze apparatus for 10 min.

*Confirmation of electrode locations by histological analysis*

The mice were overdosed with isoflurane, perfused intracardially with 4% paraformaldehyde (PFA) in phosphate-buffered saline (PBS, pH 7.4), and decapitated. After dissection, the brain was fixed overnight in 4% PFA/PBS and then cryoprotected by successive overnight incubations in 20% sucrose and 30% sucrose in PBS. Frozen coronal sections (100 μm) were cut using a microtome, mounted, and processed for cresyl violet staining. Briefly, slices were rinsed in water, stained with cresyl violet, and coverslipped with Permount. The positions of all electrodes were confirmed by identifying the corresponding electrode tracks in histological tissue sections.

**Extraction of RNA and RNA sequencing**

Total RNA was extracted from the prefrontal cortex, amygdala, and striatum of four mice per genotype (two 4-month-old and two 10-months-old individuals from batches 1 and 2), yielding 48 samples in total. Cage mates were selected for sequencing when possible (see Supplementary Fig. S3 and Table S4 for detailed sampling information). Briefly, mice were habituated to a novel cage for 30 minute and then decapitated under anesthesia. The indicated brain regions were then isolated using a mouse brain atlas (17) by the same experimenter. The extracted tissues were homogenized in BioMasher Standard (TaKaRa, Shiga, Japan) and total RNA isolated using the RNeasy® Plus Mini Kit (Qiagen, Hilden, Germany). RNA concentration and purity were assessed using a Nano-Drop® ND-1000 spectrophotometer (Thermo Scientific, Waltham, MA), and total RNA integrity quantified by the Agilent 2100 Bioanalyzer (Agilent Technologies, Santa Clara, CA). From the 48 RNA samples, 48 cDNA libraries were prepared and 100 bp paired-end reads were sequenced on a DNBSEQ platform at the Beijing Genomics Institute (BGI, Hong Kong, China). A total of 2,344,530,700 raw sequencing reads were generated (24–25 million read pairs per sample; Supplementary Table S4) and used for gene expression profiling.

**Data processing**

The raw reads were quality checked, trimmed, and filtered using fastp 0.20.0 (18), yielding 20–21 million filtered reads (Supplementary Table S4). STAR v2.7.5c (19) was then used for mapping the reads to the reference mouse genome (GRCm38.p6). We assigned reads to the Ensembl100 annotation and generated fragment counts by the featureCounts utility of the Subread package (v1.6.3) in R (20).We obtained reads with mapping quality above 20 (essentially uniquely mapped reads) and only pairs that were properly aligned on the same chromosome. The fragment counts were then used to perform differential expression analysis with iDEP v0.91 workflow (21), a web application that includes several conventional software applications such as DESeq2 (22) and edgeR (23), and can perform comprehensive analysis of RNA-Seq data. After normalizing all counts according to effective library size, we retained genes with > 1 count per million (CPM) in half of the samples (24 samples). A total of 13,608 genes were then assessed in downstream analyses. Principal component analysis (PCA) was first performed to reveal the overall expression pattern of each sample. While most samples demonstrated distinct expression patterns, one supposed amygdala sample (derived from a 4-month-old *Vmat1*$^{\mathrm{Thr/Thr}}$ mouse) exhibited an expression pattern similar to the other striatal samples (Supplementary Fig. S20). Given the inconsistency in the expression pattern, we excluded this sample and conducted the normalization step again for the remaining 47 samples.

**Differential expression and weighted gene correlation network analysis**

We conducted pair-wise between-genotype comparisons of the same brain regions to identify differentially expressed genes (DEGs), which were defined by |log2(fold change)| > 1 and false discovery rate (FDR; corrected by the Benjamini and Hochberg method) < 0.05. The DEGs were further characterized by enrichment analysis for GO terms in the iDEP pipeline. An Illumina BaseSpace application (http://basespace.illumina.com/apps) was used to investigate the correlation in expression levels of DEGs among currently and previously collected datasets. In this analysis, we set FDR < 0.05 as cutoff to select genes for input, yielding 131 genes recognized in total. To identify specific biological pathways affected by *Vmat1* genotypes, we conducted weighted gene correlation network analysis (WGCNA) (24) of 15 samples from the amygdala. First, we calculated the transcripts per million (TPM) for each gene, and the expression matrix was then adjusted for mouse batch as a potential confounding factor. The 1000 genes with most variable expression levels (largest coefficients of variation) among samples were retained. We chose a soft thresholding power = 21, as it yielded a scale-free topology fitting index (R squared) greater than 0.8. The correlations among gene expression levels across samples were then calculated to detect co-expressing modules. We also calculated the correlations between the expression levels of a given gene in the amygdala and the behavioral composite (locomotor activity and anxiety), based on the data of 8 individuals (two for each genotype). Genes belonging to modules of interest were further included in protein-protein interaction networks constructed using STRING, and functional gene clusters were detected by the MCODE module of Cytoscape 3.8.2 (25).

**Availability of data and materials**

The datasets used and/or analyzed in the current study are available from the corresponding author on reasonable request. The raw RNA-Seq reads have been deposited in the NCBI Sequence Read Archive (SRA) under BioProject accession no. PRJNA660500. Codes used for RNA-Seq analysis are available on the GitHub repository: https://github.com/daikisato12/Vmat1_RNASeq.

**Supplementary Tables and Figures**

**Supplementary Table S1. CRISPR RNAs, donor DNAs, and primers used for genome editing and genotyping**

| Aim | Name | Sequence (5' to 3') |
|---|---|---|
| Electroporation of mouse eggs | Vmat1 Exon4 crRNA | AAGAAGAAAAUGUCCGGAUUguuuuagagcuaugcuguuuug |
| | tracrRNA | AAACAGCAUAGCAAGUUAAAAUAAGGCUAGUCCGUUAUCA<br>ACUUGAAAAAGUGGCACCGAGUCGGUGCU |
| | ssDNA donor for 133Thr substitution | AAAACAACTGCTTGCAAGGGATAGAGTTCTTGGAAGAAGAAA<br>**C**TGTCCGGATTGG**A**ATTCTATTTGCTTCAAAGGCTTTGATGCA<br>ACTTCTGGTTAACCC |
| | ssDNA donor for 133Ile substitution | AAAACAACTGCTTGCAAGGGATAGAGTTCTTGGAAGAAGAAA<br>**T**TGT**G**CG**C**ATTGGGATTCTATTTGCTTCAAAGGCTTTGATGCA<br>ACTTCTGGTTAACCC |
| Genotyping (PCR-RFLP assay) | Exon4_F | GGCGGTGTCATCTTCATTAGGC |
| | Exon4_R | CCAACTCCAGAGTCATTCTTTCCC |
| Off-target analyses (OT1-OT12) | OT1_F | AGGGTTAGAAAGCAAATTTTCAGAGCC |
| | OT1_R | TCATAAACTGAAGCAGGTGGTATGTG |
| | OT2_F | TTGCTTACTACAGCCATACTTCTGAAC |
| | OT2_R | TCACCCTAAAGTAACCCAGAGTGTTC |
| | OT3_F | GTCAGAAGACCTGAAATTCTGTAGTCAC |
| | OT3_R | GCAGAGATCACTCTTACCCCTATGTTAC |
| | OT4_F | CTGGCAAAACAAAGACACTCCTATATC |

| | |
|---|---|
| OT4_R | AGGATTGAATGTGAAACACACTGGTAG |
| OT5_F | TTGCCAAAGGAGAGGGACACATGTTAC |
| OT5_R | CTGTTCTGGCTACTTTACACCTCTACTG |
| OT6_F | GAAGAATCACTGGCTGTAGATAAGTCC |
| OT6_R | GTTATATTAGTGGCTTTGGGATACAGC |
| OT7_F | TATATTGCCCTTCAATAAAACTGCATG |
| OT7_R | TACTACAACCTATACCACAAAACCGTC |
| OT8_F | AGCATGATTACCACAAACATGAAACTG |
| OT8_R | AGTAGATCAGATGGCTCCTTGGTATC |
| OT9_F | AGGACTTTGCTTACAAAACAGAAAGC |
| OT9_R | TTACCCAGCTCACTATATGTTCCCTAC |
| OT10_F | GACCTCCACAAATTACCATGACATAC |
| OT10_R | CTTGTGTGTATGTAAGCAGAGAGCAG |
| OT11_F | TAACTGGGCTAAAAGATGGCTTAGTG |
| OT11_R | AGTGATTCTCCATGCTTTTTGTAGTG |
| OT12_F | AAGTAGTTCCCATTTTACACGAGAGTC |
| OT12_R | GACCAAACACTGTAGACTTGAGGTTC |

*Bold; intended substitutions, underlined; homology arms

**Supplementary Table S2. Potential off-target loci (OT1-OT12) for the *Vmat1* exon 4 guide RNA sequence as predicted by the CRISPOR web-tool**

| Name | Off-target Sequence* | Mismatch position | Mismatch count | MIT Off-target Score | CFD Off-target Score | Chr | Start | End | Strand | Locus Description |
|---|---|---|---|---|---|---|---|---|---|---|
| OT1 | AAGAAGAAAATGGCTGGATTGGG | ............*.*..... | 2 | 0.566 | 0.017 | 2 | 16686819 | 16686841 | - | intron:Plxdc2 |
| OT2 | AAGGAAAAGCTGTCCGGATTAGG | ...*.*..**.......... | 4 | 0.465 | 0.139 | 13 | 106559519 | 106559541 | + | intergenic:Dph3b-ps-Gm24527 |
| OT3 | AAGGACAAAATGTCTGGATTTGG | ...*.*........*..... | 3 | 0.456 | 0.028 | 1 | 190717633 | 190717655 | + | intron:Rps6kc1 |
| OT4 | TAGTAGGAAATGTCCGTATTTGG | *..*..*.........*... | 4 | 0.416 | 0.112 | 4 | 27570801 | 27570823 | - | intergenic:Gm11901-Gm25173 |
| OT5 | ACGAAAAAATTGTCCAGATTAGG | .*...*...*.....*.... | 4 | 0.144 | 0.693 | 2 | 60849507 | 60849529 | + | intron:Rbms1 |
| OT6 | TAGAAGGAAATGCCCAGATTTGG | *.....*.....*..*.... | 4 | 0.072 | 0.557 | 12 | 65526603 | 65526625 | - | intergenic:Gm26015-Gm25599 |
| OT7 | AAGAAAAAAATCTCCAAATTGGG | .....*.....*...**... | 4 | 0.028 | 0.494 | 6 | 45559080 | 45559102 | - | intron:Cntnap2 |

| | | | | | | | | | | |
|---|---|---|---|---|---|---|---|---|---|---|
| OT8 | AAGAATAAAATATTCAGATTGGG | .....*.....*.*.*.... | 4 | 0.011 | 0.456 | 8 | 89723230 | 89723252 | + | intergenic:Gm24212-Tox3 |
| OT9 | AAAAAGAAAAAGTTCAGATTAGG | ..*.......*..*.*.... | 4 | 0.021 | 0.412 | 7 | 139388181 | 139388203 | - | intergenic:Pwwp2b-Inpp5a |
| OT10 | TAGAATAAAAAGCCCGGATTGGG | *....*....*.*....... | 4 | 0.195 | 0.395 | 2 | 105242794 | 105242816 | + | intron:0610012H03Rik |
| OT11 | AAAAAGAAAAAGACCAGATTAGG | ..*.......*.*..*.... | 4 | 0.055 | 0.389 | 8 | 111294818 | 111294840 | + | intron:Rfwd3 |
| OT12 | AAGAAGAACTTGTTCGAATTCGG | ........**...*..*... | 4 | 0.044 | 0.345 | 17 | 55701468 | 55701490 | + | intron:Pot1b |

*On target sequence: AAGAAGAAAATGTCCGGATTGGG (chr8:69073624-69073646, mm10)

**Supplementary Table S3. Characteristics of mice in behavioral tests**

| Experiment | The number of individuals tested | Age (weeks)* |
|---|---|---|
| General health and neurological screening | WT: 20, Thr/Thr: 20, Thr/Ile: 20, Ile/Ile: 20 | 9–12 |
| Light/Dark transition | WT: 20, Thr/Thr: 20, Thr/Ile: 20, Ile/Ile: 20 | 10–13 |
| Open field | WT: 20, Thr/Thr: 20, Thr/Ile: 20, Ile/Ile: 20 | 10–13 |
| Elevated plus maze | WT: 20, Thr/Thr: 20, Thr/Ile: 20, Ile/Ile: 20 | 11–14 |
| Hot plate | WT: 20, Thr/Thr: 20, Thr/Ile: 20, Ile/Ile: 20 | 11–14 |
| Social interaction in a novel environment | WT: 20, Thr/Thr: 20, Thr/Ile: 20, Ile/Ile: 20 | 12–15 |
| Rotarod | WT: 20, Thr/Thr: 20, Thr/Ile: 20, Ile/Ile: 20 | 12–16 |
| Three-chambers social interaction | WT: 20, Thr/Thr: 20, Thr/Ile: 20, Ile/Ile: 20 | 15–19 |
| Startle response / prepulse inhibition | WT: 20, Thr/Thr: 20, Thr/Ile: 20, Ile/Ile: 20 | 16–19 |
| Porsolt forced swim | WT: 20, Thr/Thr: 19, Thr/Ile: 20, Ile/Ile: 20 | 16–19 |
| T-maze | WT: 20, Thr/Thr: 19, Thr/Ile: 20, Ile/Ile: 20 | 18–22 |
| Bars maze | WT: 20, Thr/Thr: 19, Thr/Ile: 20, Ile/Ile: 20 | 19–30 |
| Tail suspension | WT: 20, Thr/Thr: 19, Thr/Ile: 20, Ile/Ile: 20 | 28–31 |
| Cued and contextual fear conditioning | WT: 20, Thr/Thr: 18, Thr/Ile: 19, Ile/Ile: 20 | 28–35 |
| Home cage social interaction | WT: 20, Thr/Thr: 18, Thr/Ile: 18, Ile/Ile: 20 | 35–39 |
| Impulsivity | Thr/Thr: 10, Ile/Ile: 10 | 37–55 |

*Age (weeks) is shown in weeks at the beginning of each test.

**Supplementary Table S4. Summary of samples and reads for transcriptome analysis**

| ID | Genotype | Age (month) | Tissue | Raw reads (pairs) | Filtered reads | % out of raw reads | Uniquely mapped reads | % out of filtered reads | RIN |
|---|---|---|---|---|---|---|---|---|---|
| 10P | Thr/Thr | 10 | Prefrontal cortex | 24,383,779 | 20,859,057 | 85.5 | 18,993,167 | 91.1 | 9.2 |
| 10A | Thr/Thr | 10 | Amygdala | 24,087,830 | 20,921,381 | 86.9 | 18,880,735 | 90.2 | 9.2 |
| 10S | Thr/Thr | 10 | Striatum | 24,115,209 | 20,654,411 | 85.6 | 18,431,340 | 89.2 | 9.2 |
| 12P | WT | 10 | Prefrontal cortex | 24,092,820 | 20,735,622 | 86.1 | 18,737,622 | 90.4 | 8.8 |
| 12A | WT | 10 | Amygdala | 24,081,462 | 20,840,090 | 86.5 | 18,911,739 | 90.7 | 8.7 |
| 12S | WT | 10 | Striatum | 24,035,315 | 20,530,610 | 85.4 | 18,472,395 | 90.0 | 9.4 |
| 13P | WT | 10 | Prefrontal cortex | 24,143,724 | 20,729,730 | 85.9 | 18,677,477 | 90.1 | 8.8 |
| 13A | WT | 10 | Amygdala | 24,122,229 | 21,000,435 | 87.1 | 18,815,021 | 89.6 | 8.8 |
| 13S | WT | 10 | Striatum | 24,106,695 | 20,387,696 | 84.6 | 18,227,770 | 89.4 | 9.5 |
| 14P | Thr/Thr | 10 | Prefrontal cortex | 24,087,074 | 20,576,341 | 85.4 | 18,577,500 | 90.3 | 9.0 |
| 14A | Thr/Thr | 10 | Amygdala | 24,157,397 | 20,810,909 | 86.1 | 18,884,533 | 90.7 | 9.0 |
| 14S | Thr/Thr | 10 | Striatum | 24,229,830 | 21,155,028 | 87.3 | 18,992,814 | 89.8 | 9.4 |
| 15P | Thr/Ile | 10 | Prefrontal cortex | 24,141,270 | 21,011,593 | 87.0 | 19,153,852 | 91.2 | 9.4 |
| 15A | Thr/Ile | 10 | Amygdala | 24,094,205 | 20,465,526 | 84.9 | 18,554,809 | 90.7 | 9.3 |
| 15S | Thr/Ile | 10 | Striatum | 24,110,797 | 20,124,957 | 83.5 | 18,211,642 | 90.5 | 9.5 |
| 16P | Ile/Ile | 10 | Prefrontal cortex | 24,096,840 | 19,998,181 | 83.0 | 18,378,018 | 91.9 | 9.4 |
| 16A | Ile/Ile | 10 | Amygdala | 24,085,769 | 20,050,290 | 83.2 | 17,968,973 | 89.6 | 9.1 |

| | | | | | | | | | |
|---|---|---|---|---|---|---|---|---|---|
| 16S | Ile/Ile | 10 | Striatum | 24,224,639 | 20,792,275 | 85.8 | 18,500,603 | 89.0 | 9.3 |
| 17P | Thr/Thr | 4 | Prefrontal cortex | 24,053,586 | 20,114,080 | 83.6 | 18,292,384 | 90.9 | 9.0 |
| 17A | Thr/Thr | 4 | Amygdala | 24,105,790 | 20,307,170 | 84.2 | 18,190,848 | 89.6 | 9.3 |
| 17S | Thr/Thr | 4 | Striatum | 24,184,323 | 20,366,027 | 84.2 | 18,320,470 | 90.0 | 9.3 |
| 18P | Ile/Ile | 4 | Prefrontal cortex | 24,175,686 | 20,485,560 | 84.7 | 18,559,296 | 90.6 | 8.6 |
| 18A | Ile/Ile | 4 | Amygdala | 25,119,422 | 20,799,894 | 82.8 | 18,281,497 | 87.9 | 8.1 |
| 18S | Ile/Ile | 4 | Striatum | 24,255,146 | 20,507,485 | 84.5 | 18,112,396 | 88.3 | 9.2 |
| 19P | Thr/Ile | 4 | Prefrontal cortex | 24,155,485 | 20,127,544 | 83.3 | 18,217,183 | 90.5 | 9.0 |
| 19A | Thr/Ile | 4 | Amygdala | 24,913,488 | 21,148,108 | 84.9 | 18,902,320 | 89.4 | 9.1 |
| 19S | Thr/Ile | 4 | Striatum | 24,938,317 | 20,622,885 | 82.7 | 18,170,276 | 88.1 | 8.4 |
| 20P | WT | 4 | Prefrontal cortex | 25,114,676 | 21,750,542 | 86.6 | 19,508,651 | 89.7 | 8.4 |
| 20A | WT | 4 | Amygdala | 24,925,216 | 20,813,471 | 83.5 | 18,454,605 | 88.7 | 9.0 |
| 20S | WT | 4 | Striatum | 25,017,047 | 21,220,021 | 84.8 | 18,720,098 | 88.2 | 8.5 |
| 21P | Thr/Thr | 4 | Prefrontal cortex | 24,865,805 | 20,783,104 | 83.6 | 18,891,783 | 90.9 | 9.1 |
| 21A | Thr/Thr | 4 | Amygdala | 24,968,548 | 21,272,091 | 85.2 | 19,372,403 | 91.1 | 9.1 |
| 21S | Thr/Thr | 4 | Striatum | 24,987,027 | 21,040,617 | 84.2 | 18,697,540 | 88.9 | 8.8 |
| 22P | Ile/Ile | 4 | Prefrontal cortex | 25,108,881 | 21,533,868 | 85.8 | 19,453,865 | 90.3 | 9.1 |
| 22A | Ile/Ile | 4 | Amygdala | 24,935,536 | 20,994,619 | 84.2 | 18,751,867 | 89.3 | 8.8 |
| 22S | Ile/Ile | 4 | Striatum | 24,838,609 | 20,729,654 | 83.5 | 18,423,389 | 88.9 | 9.2 |
| 23P | Thr/Ile | 4 | Prefrontal cortex | 25,048,439 | 21,639,401 | 86.4 | 19,558,627 | 90.4 | 9.2 |
| 23A | Thr/Ile | 4 | Amygdala | 24,215,091 | 20,289,031 | 83.8 | 18,105,056 | 89.2 | 9.3 |

| | | | | | | | | | |
|---|---|---|---|---|---|---|---|---|---|
| 23S | Thr/Ile | 4 | Striatum | 24,446,249 | 20,629,405 | 84.4 | 18,295,304 | 88.7 | 9.1 |
| 24P | WT | 4 | Prefrontal cortex | 24,371,757 | 20,541,360 | 84.3 | 18,556,300 | 90.3 | 9.1 |
| 24A | WT | 4 | Amygdala | 24,394,598 | 20,586,087 | 84.4 | 18,646,645 | 90.6 | 9.2 |
| 24S | WT | 4 | Striatum | 24,500,916 | 20,777,569 | 84.8 | 18,360,727 | 88.4 | 9.2 |
| 25P | Ile/Ile | 10 | Prefrontal cortex | 24,343,251 | 20,534,776 | 84.4 | 18,780,504 | 91.5 | 9.1 |
| 25A | Ile/Ile | 10 | Amygdala | 24,349,843 | 20,914,988 | 85.9 | 18,892,387 | 90.3 | 8.4 |
| 25S | Ile/Ile | 10 | Striatum | 24,434,301 | 20,522,945 | 84.0 | 18,096,483 | 88.2 | 9.0 |
| 26P | Thr/Ile | 10 | Prefrontal cortex | 24,364,878 | 20,626,166 | 84.7 | 18,730,292 | 90.8 | 9.0 |
| 26A | Thr/Ile | 10 | Amygdala | 24,295,041 | 20,266,521 | 83.4 | 18,307,648 | 90.3 | 8.6 |
| 26S | Thr/Ile | 10 | Striatum | 24,441,514 | 20,702,900 | 84.7 | 18,437,771 | 89.1 | 9.2 |

**Supplementary Table S5. Top 10 datasets from 6 independent studies showing DEGs significantly overlapped with the present study (WT vs. Ile comparison) obtained from BaseSpace analysis.** Texts in bold indicate studies of interest (related to Huntington disease) further investigated in Supplementary Figure S23.

| Study | Accession ID | Dataset | Correlation direction | Common genes | *P* value |
|---|---|---|---|---|---|
| **Study 1: Cortical and striatal tissues from R6/2 transgenic mouse model of Huntington disease** | **GSE48962** | **Striata of 12wk old R6/2 mice expressing CAG repeat expansion of huHTT gene vs wildtype** | **+** | **70** | **$2.3 \times 10^{-36}$** |
| Study 2: Cerebella and striata of SNCA KO and wildtype mice at different timepoints | GSE19534 | Neural tissues from wildtype 6mo post-natal mice - striata vs cerebella | – | 103 | $2.0 \times 10^{-32}$ |
| Study 2: Cerebella and striata of SNCA KO and wildtype mice at different timepoints | GSE19534 | Neural tissues from SNCA KO 6mo post-natal mice - striata vs cerebella | – | 108 | $1.5 \times 10^{-30}$ |
| Study 3: CNS cell types - Drd1 and Drd2 Medium spiny neurons, motor, and Purkinje neurons | GSE13394 | Medium spiny neurons from mouse striatum - Drd1a vs whole brain lacking striatum | – | 104 | $1.5 \times 10^{-30}$ |
| Study 3: CNS cell types - Drd1 and Drd2 Medium spiny neurons, motor, and Purkinje neurons | GSE13394 | Medium spiny neurons from mouse striatum - Drd2 vs whole brain lacking striatum | – | 109 | $1.6 \times 10^{-30}$ |
| Study 4: Brain striata from Huntington disease model mice treated with HSP90 inhibitor (HSP990) | GSE29681 | Striata of 12wk old 4hr post 12mg/kg NVP-HSP990 - R6/2 mice expressing mutant Htt vs wildtype mice | + | 95 | $3.7 \times 10^{-30}$ |

| | | | | | |
|---|---|---|---|---|---|
| **Study 4: Brain striata from Huntington disease model mice treated with HSP90 inhibitor (HSP990)** | **GSE29681** | **Striata of 12wk old 4hr post vehicle - R6/2 mice expressing mutant Htt vs wildtype mice** | + | **88** | $\mathbf{5.5 \times 10^{-30}}$ |
| Study 2: Cerebella and striata of SNCA KO and wildtype mice at different timepoints | GSE19534 | Neural tissues from wildtype 21mo post-natal mice - striata vs cerebella | – | 110 | $8.4 \times 10^{-30}$ |
| **Study 5: Striata at 8wk from Huntington disease mouse models with or without P62 knockout** | **GSE62210** | **Striata at 8wk from P62 knockout mice – Huntington' s disease models vs wildtype** | + | **66** | $\mathbf{3.4 \times 10^{-29}}$ |
| Study 6: Striatum and cerebellum expression in Huntington disease Hdh mutant mice | GSE19780 | Brain of Huntington's disease Hdh mutant (Q111) CD1 3-10wk old mice - striatum vs cerebellum | – | 85 | $5.1 \times 10^{-29}$ |

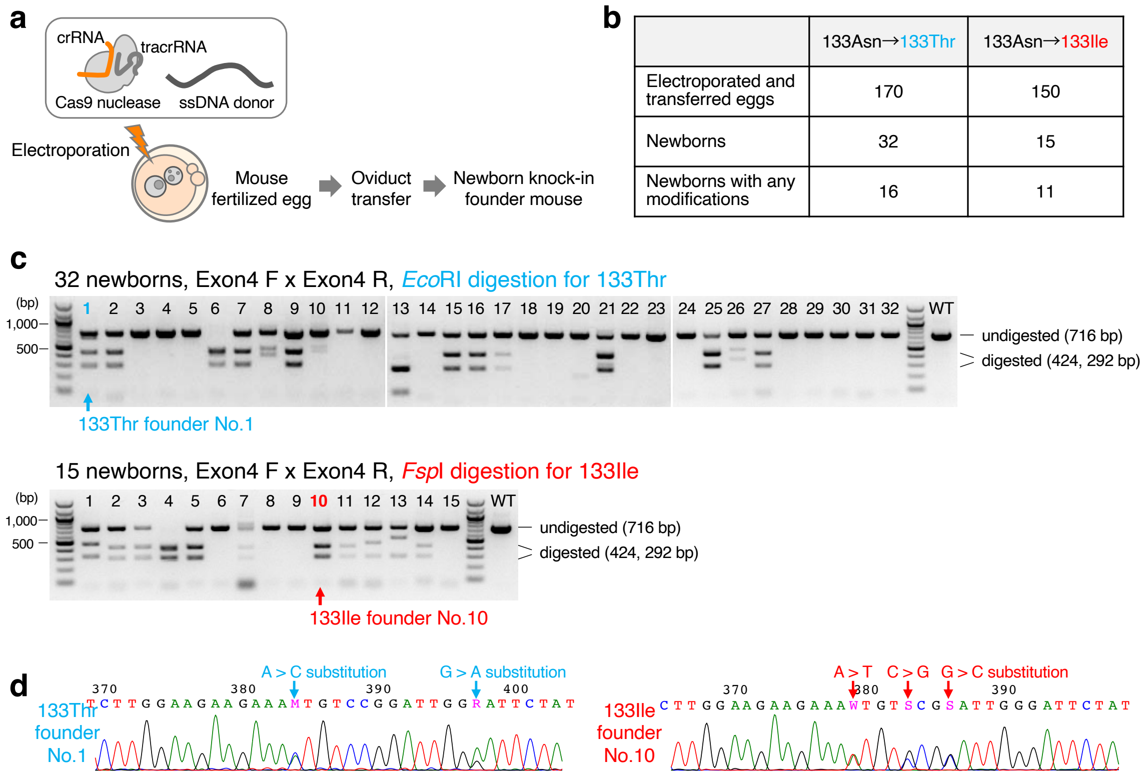


**Supplementary Figure S1. Generation of the Vmat1-humanized mouse models by one-cell zygote electroporation. (a)** Schematic diagram of electroporation-mediated CRISPR/Cas9 genome editing. Two parts of the CRISPR guide RNA (tracrRNA and crRNA) for *Vmat1* exon 4 (listed in Supplementary Table S1), recombinant Cas9 nuclease, and ssDNA donors containing the intended substitutions were electroporated into C57BL/6J mouse fertilized eggs. The eggs were subsequently transferred to surrogate mothers to obtain founder knock-in mice. **(b)** and **(c)** Results of electroporation-mediated genome editing in mouse eggs. Newborns were screened by PCR-RFLP assay as shown in Fig. 2b. PCR products were cleaved only in the presence of restriction enzyme recognition sites (*Eco*RI for 133Thr, *Fsp*I for 133Ile). CRISPR/Cas9-mediated gene modification was highly efficient. More than 50% of newborns harbored knock-in alleles, although they could have inaccurate editing. We analyzed all knock-in candidates by sequencing, which confirmed that 133Thr No.1 and 133Ile No.10 carried the correct substitutions as designed (see Fig. 2a). **(d)** Sanger sequencing results from 133Thr founder No.1 and 133Ile founder No.10 are summarized.

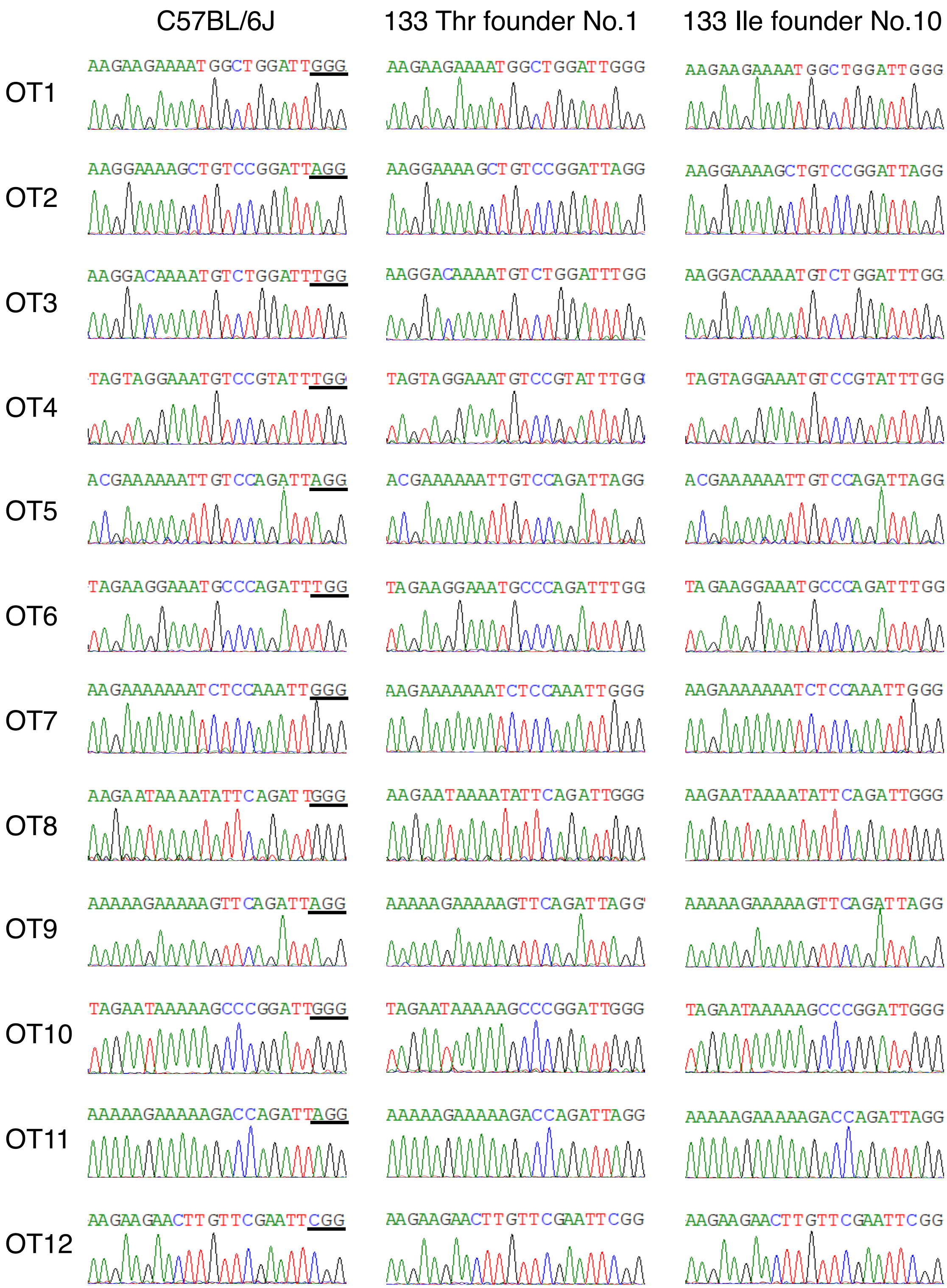


**Supplementary Figure S2. Sequencing results for 12 candidate off-target loci in founder mice (133Thr No.1, 133Ile No.10) and a wild type control mouse.** No signs of off-target cleavages were detected within the genomic regions evaluated. The PAM sequences are underlined for the 12 candidate loci.

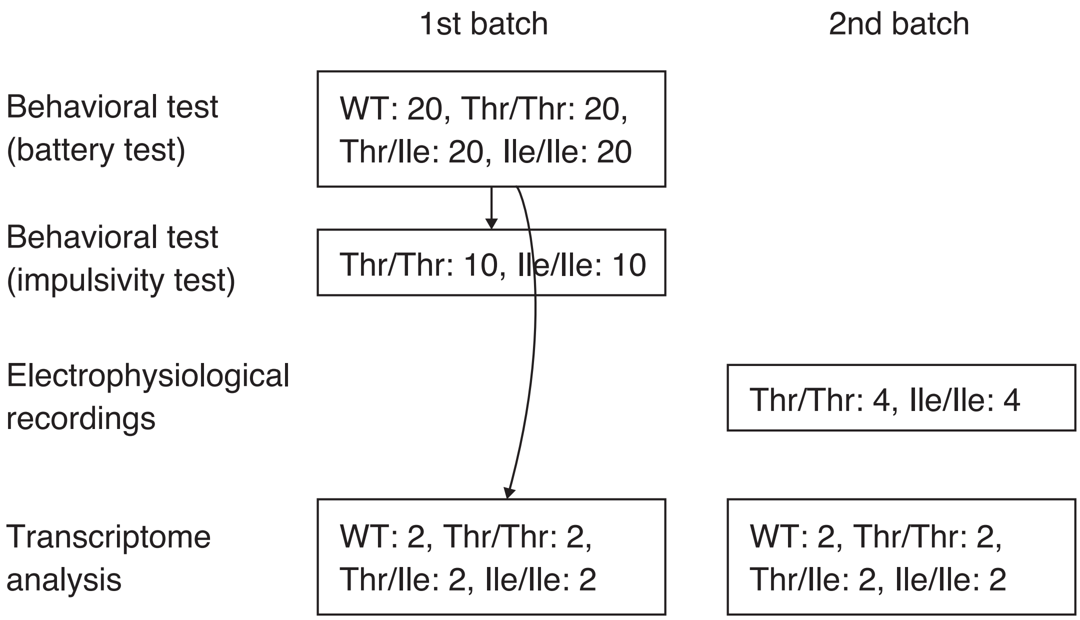


**Supplementary Figure S3. Samples and batch information for each experiment and analysis.**

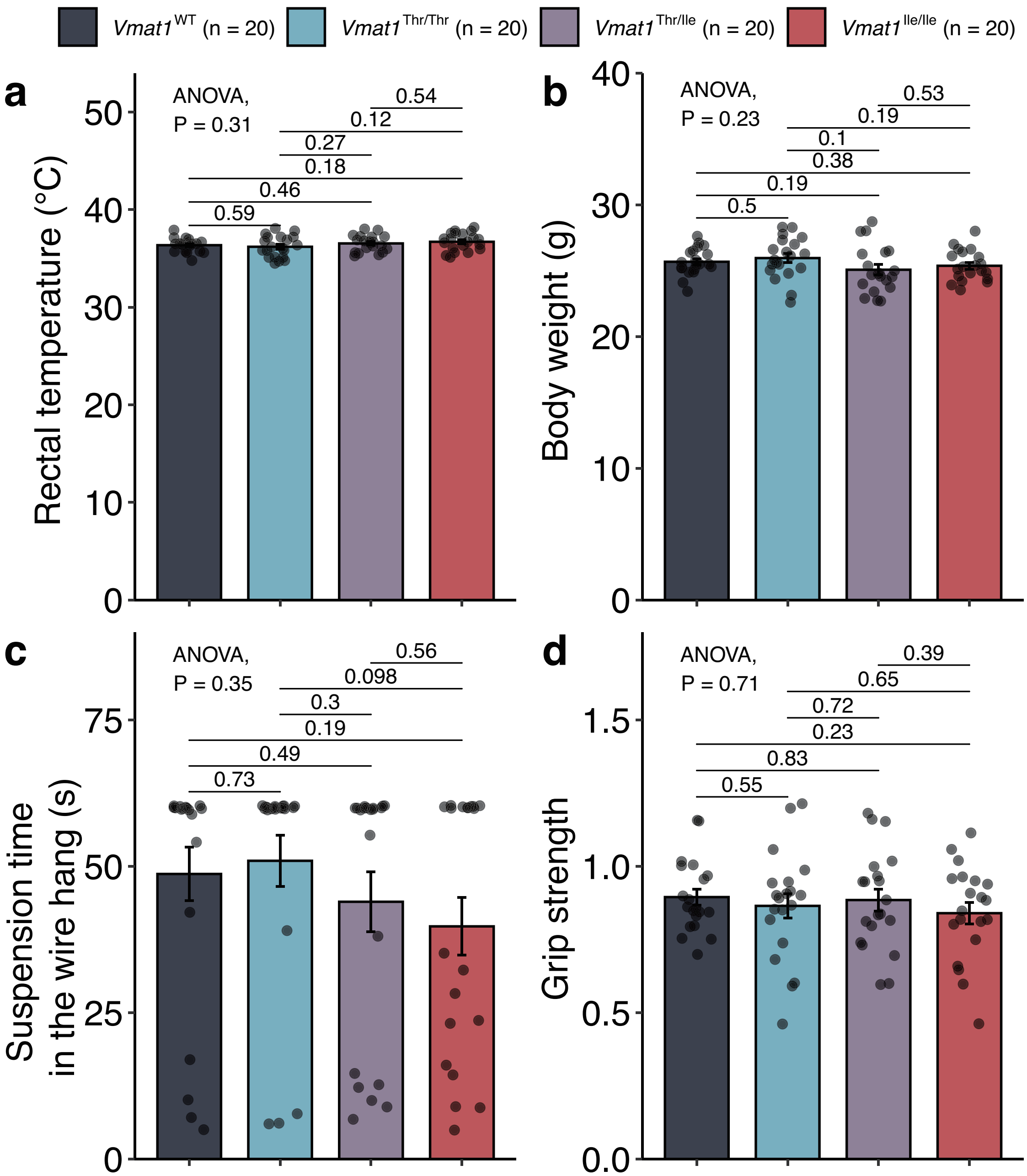


**Supplementary Figure S4. General health and neurological screening results. (a)** Rectal temperature, **(b)** body weight, **(c)** suspension time in the wire hanging test, and **(d)** grip strength. *P* values were calculated by one-way ANOVA and pair-wise *t*-tests (uncorrected). Error bars represent standard errors.

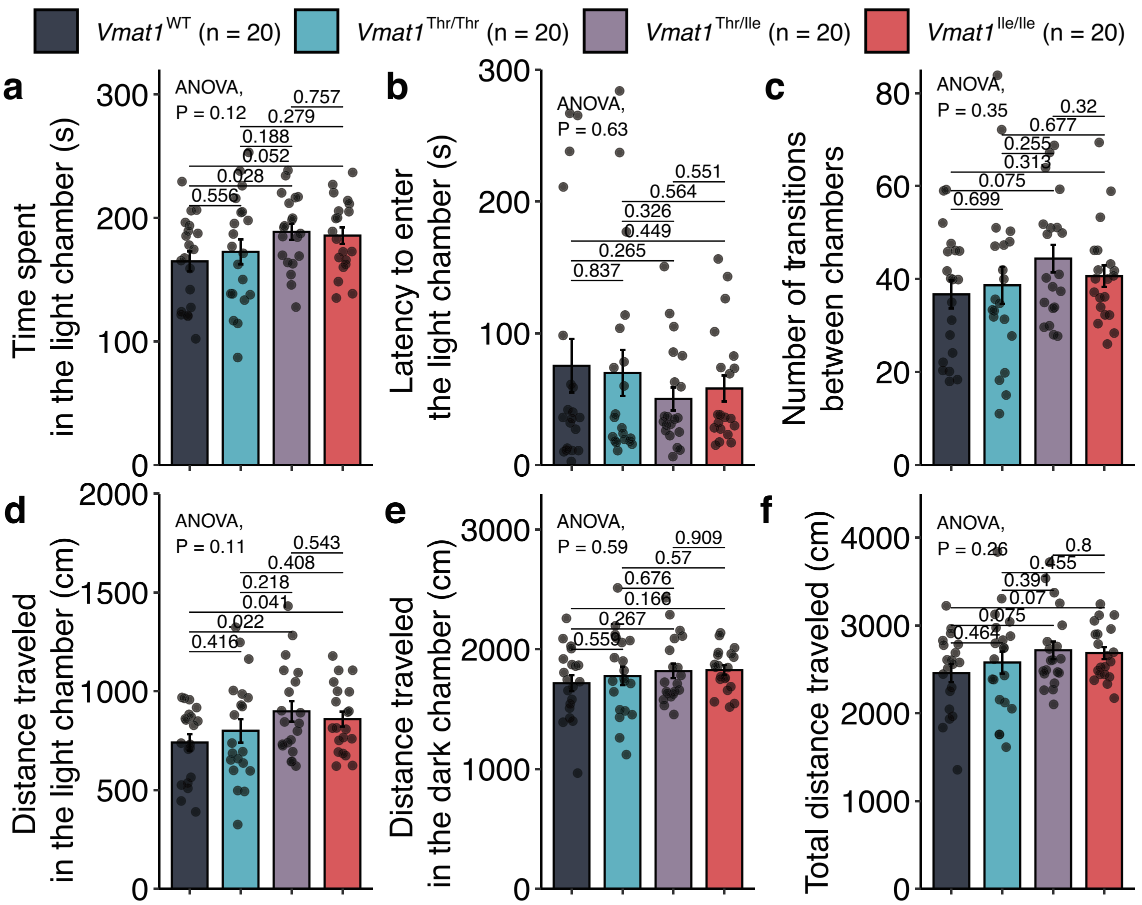


**Supplementary Figure S5. Mouse movement during the light/dark transition test. (a)** Time spent in the light chamber, **(b)** latency to enter the light chamber, **(c)** number of transitions between chambers, **(d)** distance traveled in the light chamber, **(e)** distance traveled in the dark chamber, and **(f)** total distance traveled. *P* values were calculated by one-way ANOVA and pair-wise *t*-tests (uncorrected). Error bars represent standard errors.

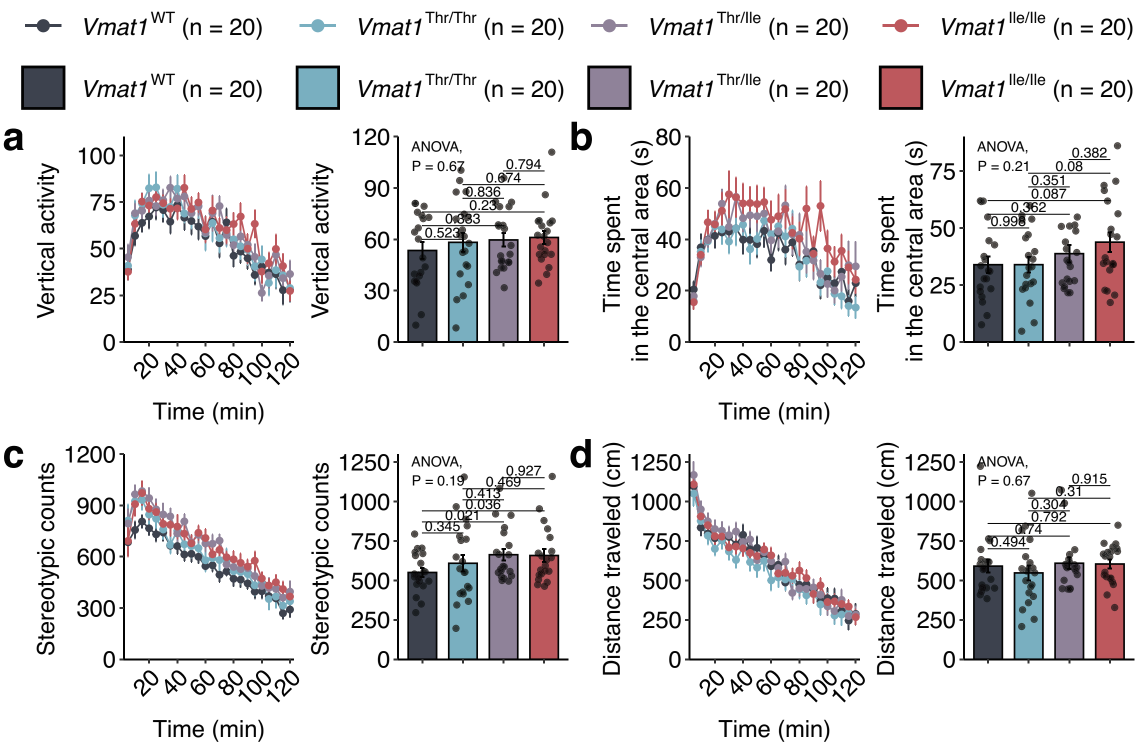


**Supplementary Figure S6. Summary of open field test behaviors. (a)** Vertical activity, **(b)** time spent in the central area, **(c)** number of stereotypic behaviors, and **(d)** total distance traveled. For each behavioral parameter, the left panels plot values over time and the right panels plot the average throughout the 2-h test. *P* values were calculated by one-way ANOVA and pair-wise *t*-tests (uncorrected). Error bars represent standard errors.

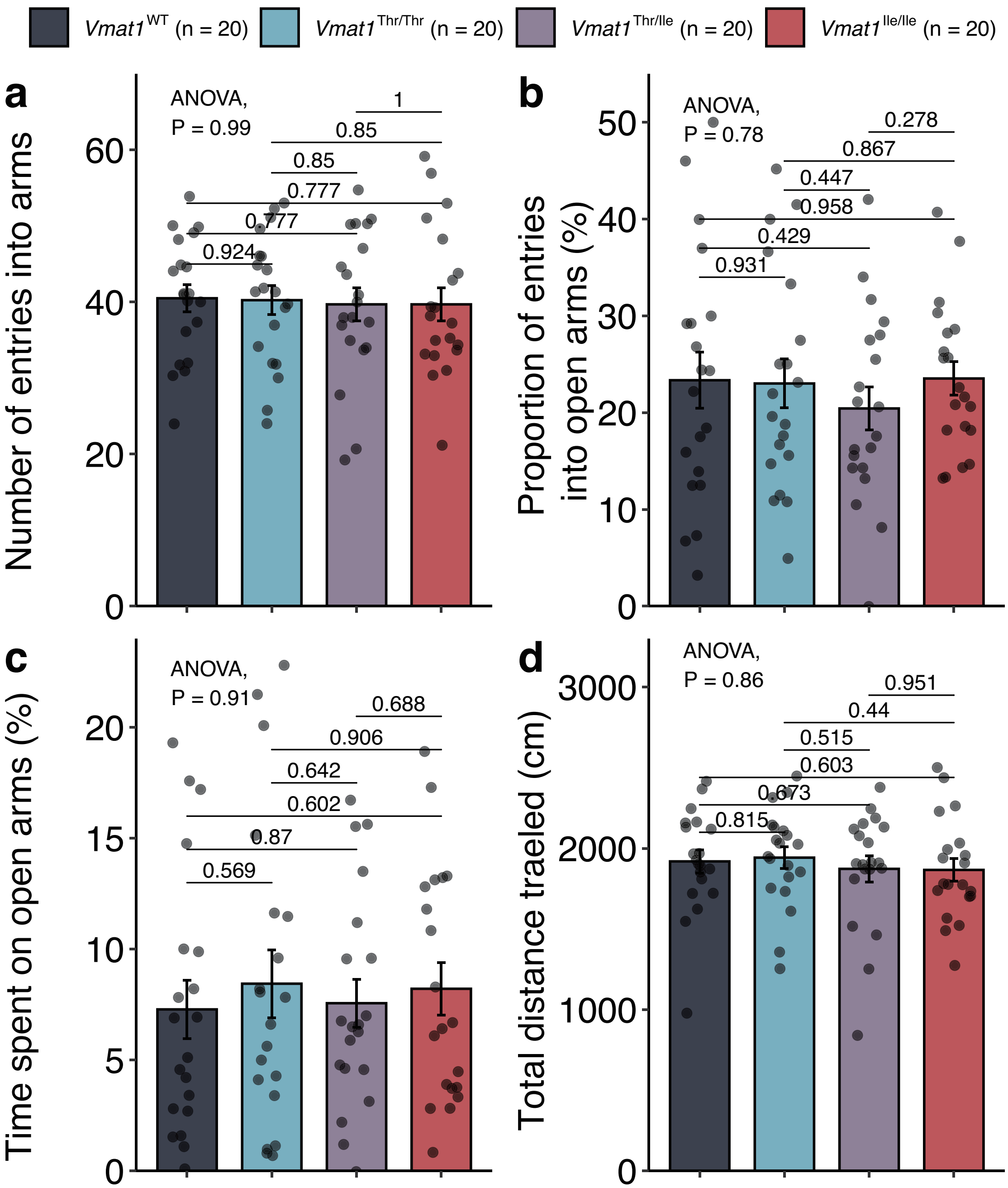


**Supplementary Figure S7. Summary of behavioral parameters during the elevated plus maze test. (a)** Number of entries into arms (open+closed), **(b)** proportion (%) of entries into open arms, **(c)** time spent on open arms, and **(d)** total distance traveled. *P* values were calculated by one-way ANOVA and pair-wise *t*-tests (uncorrected). Error bars represent standard errors.

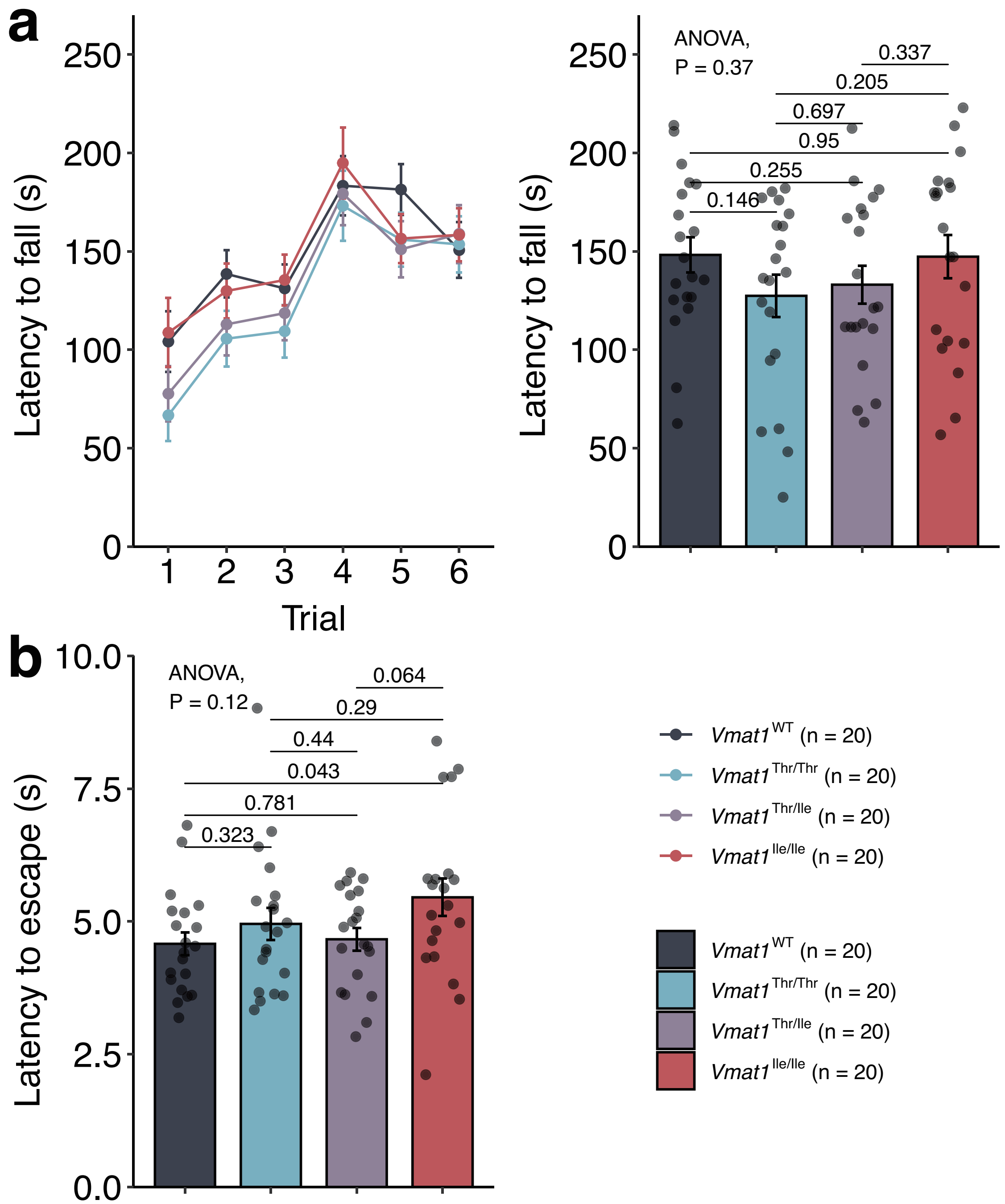


**Supplementary Figure S8. Rotarod and hot plate test results. (a)** Latency to fall from the rotarod. The left panel plots the average time to fall by trial number and the right panel plots the overall average across trials. **(b)** Latency to escape from a hot plate. *P* values were calculated by one-way ANOVA and pair-wise *t*-tests (uncorrected). Error bars represent standard errors.

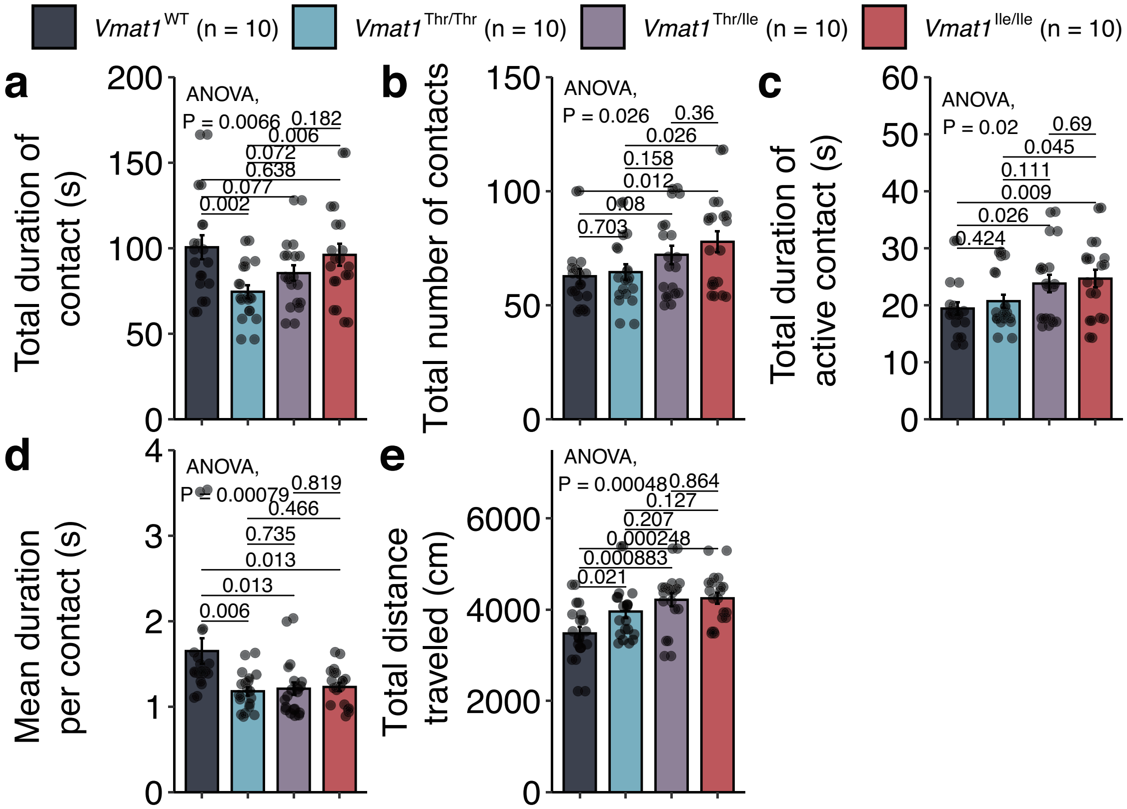


**Supplementary Figure S9. Single-chamber social interaction test results. (a)** Total duration of contact, **(b)** total number of contacts, **(c)** total duration of active contact, **(d)** mean duration per contact, and **(e)** total distance traveled. *P* values were calculated by one-way ANOVA and pair-wise *t*-tests (uncorrected). Error bars represent standard errors.

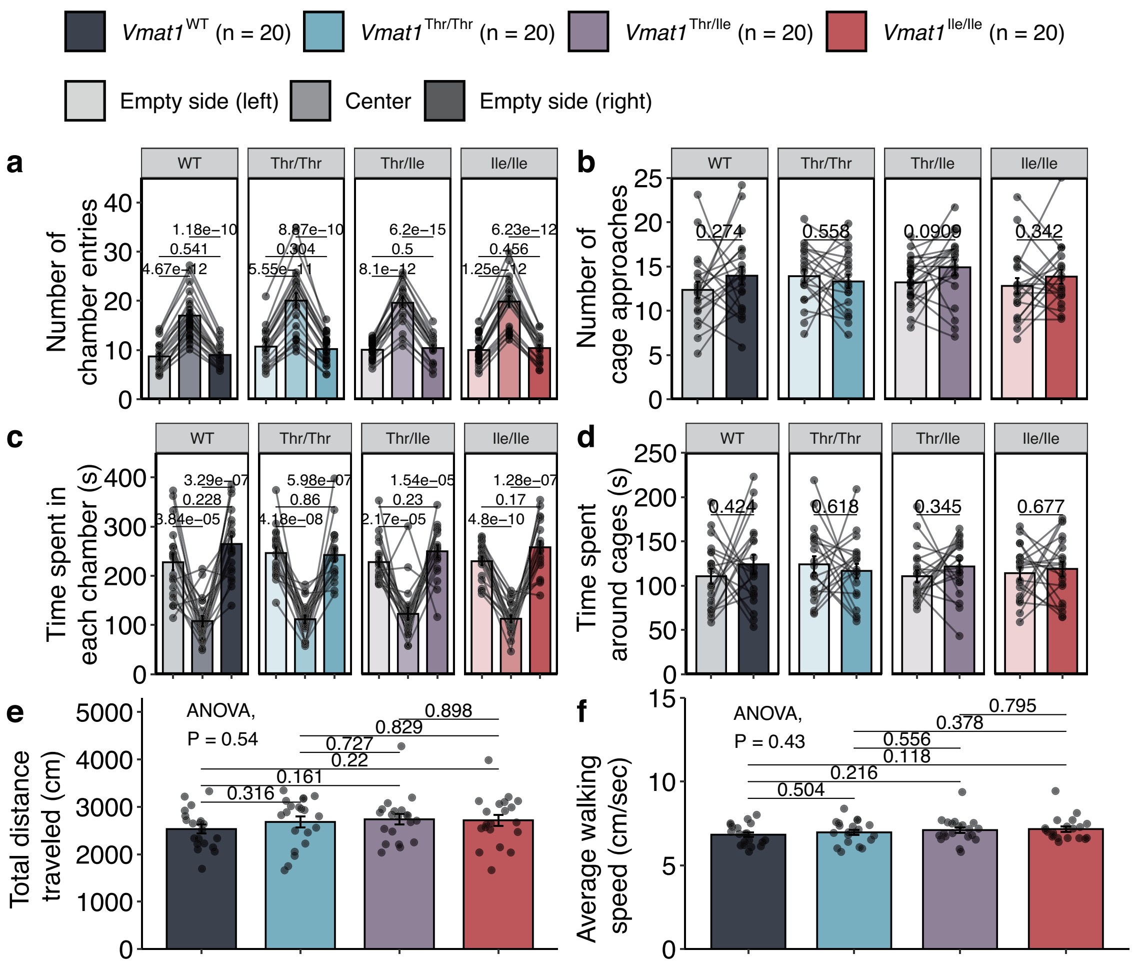


**Supplementary Figure S10-1. Crawley's 3-chamber social interaction test results (habituation phase). (a)** Number of chamber entries, **(b)** number of cage approaches, **(c)** time spent in each chamber, **(d)** time spent around cages, **(e)** total distance traveled, and **(f)** average walking speed during the test. *P* values were calculated by paired *t*-tests (uncorrected; for **(a)**–**(d)**) and one-way ANOVA (**(e) and (f)**). Error bars represent standard errors.

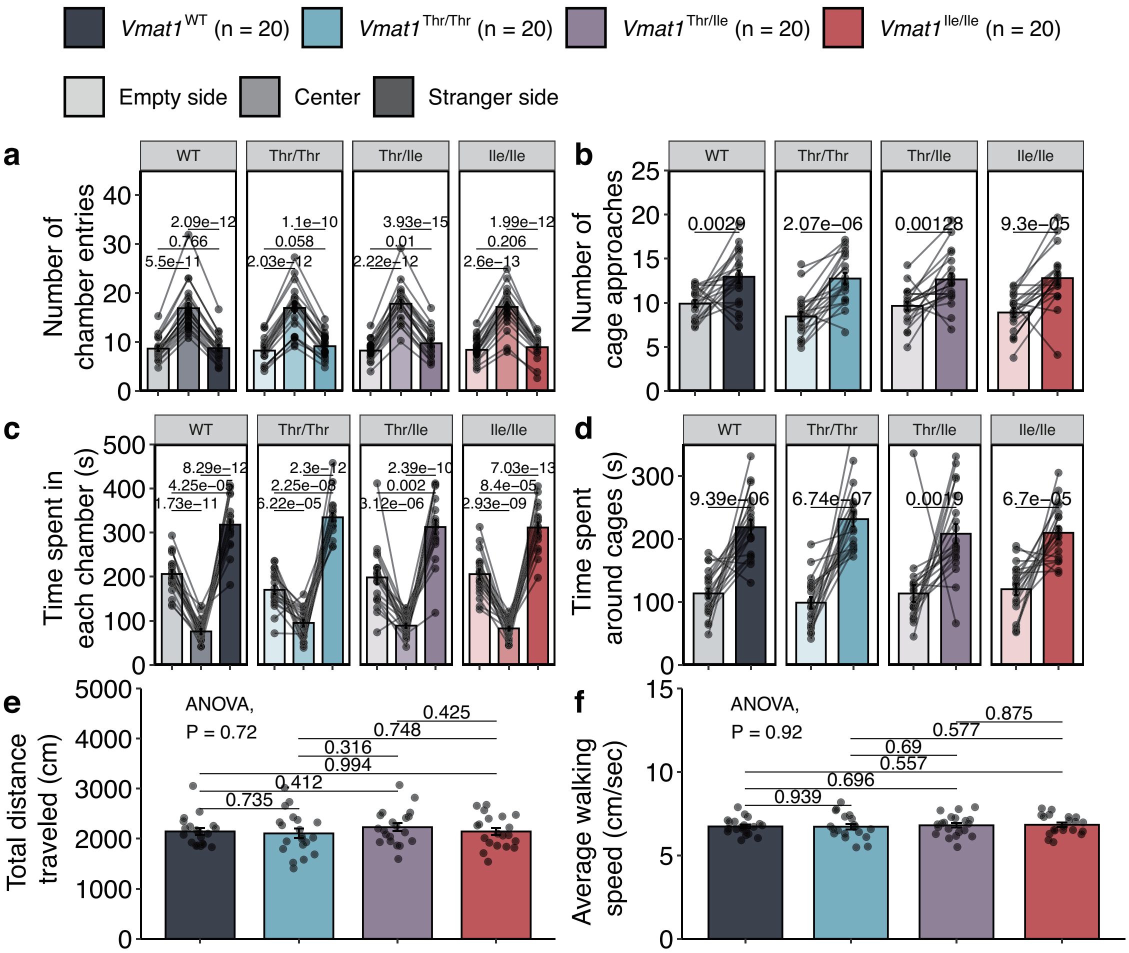


**Supplementary Figure S10-2. Crawley's 3-chamber social interaction test results (sociality). (a)** Number of chamber entries, **(b)** number of cage approaches, **(c)** time spent in each chamber, **(d)** time spent around cages, **(e)** total distance traveled, and **(f)** average walking speed during the test. *P* values were calculated by paired *t*-tests (uncorrected; for **(a)–(d)**) and one-way ANOVA (**(e) and (f)**). Error bars represent standard errors.

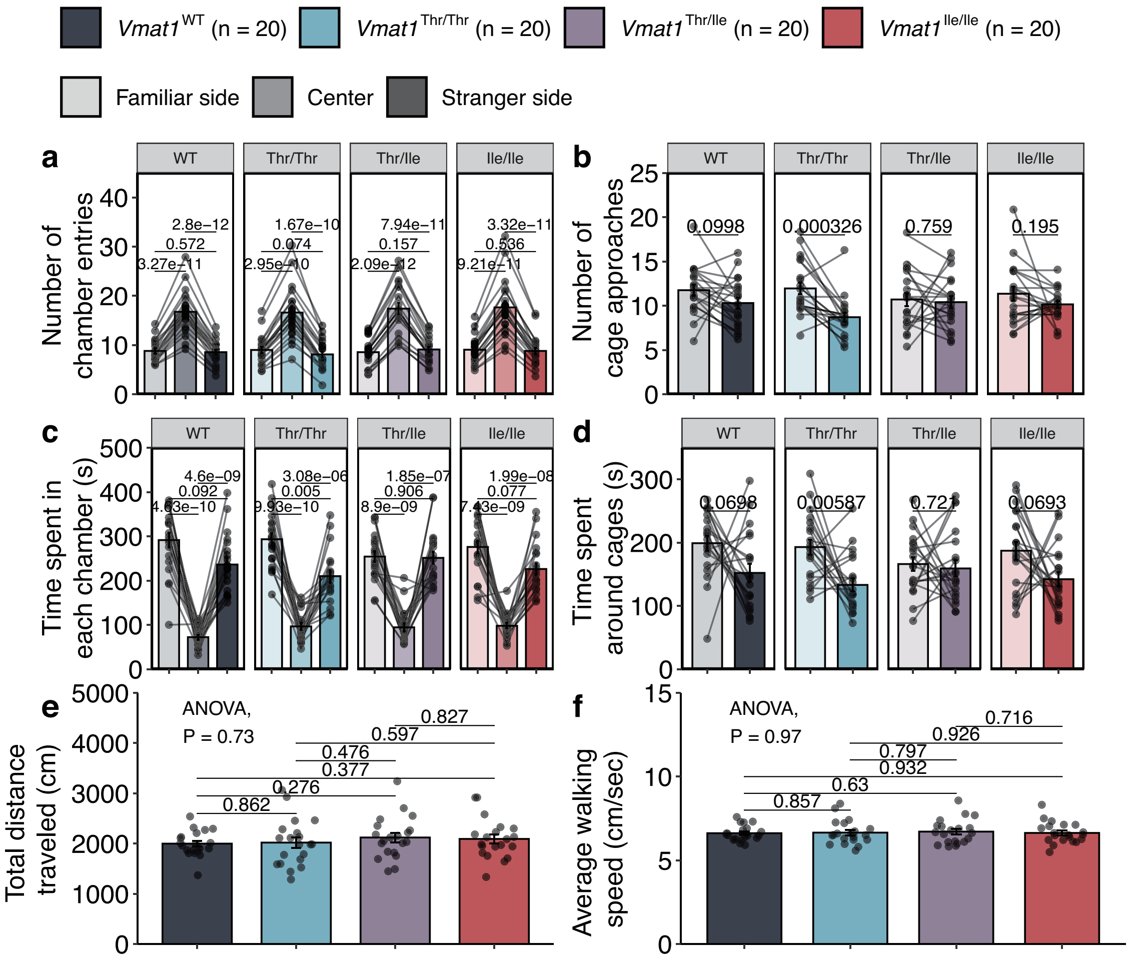


**Supplementary Figure S10-3. Crawley's 3-chamber social interaction test results (preference).** **(a)** Number of chamber entries, **(b)** Number of cage approaches, **(c)** time spent in each chamber, **(d)** time spent around cages, **(e)** total distance traveled, and **(f)** average walking speed during the test. *P* values were calculated by paired *t*-tests (uncorrected; for **(a)–(d)**) and one-way ANOVA (**(e) and (f)**). Error bars represent standard errors.

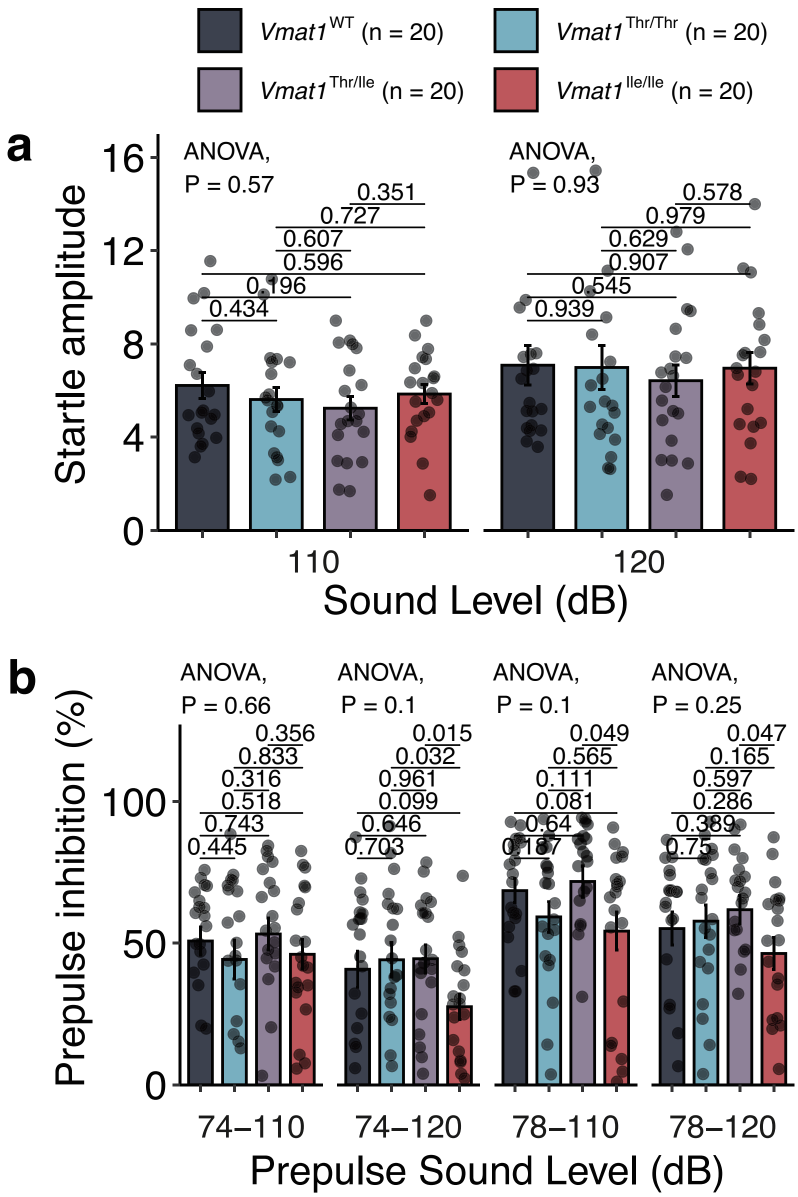


**Supplementary Figure S11. Startle responses and prepulse inhibition. (a)** Startle amplitudes in response to the loud pulse alone and **(b)** prepulse inhibition under different trial conditions (prepulse intensities). Error bars represent standard errors.

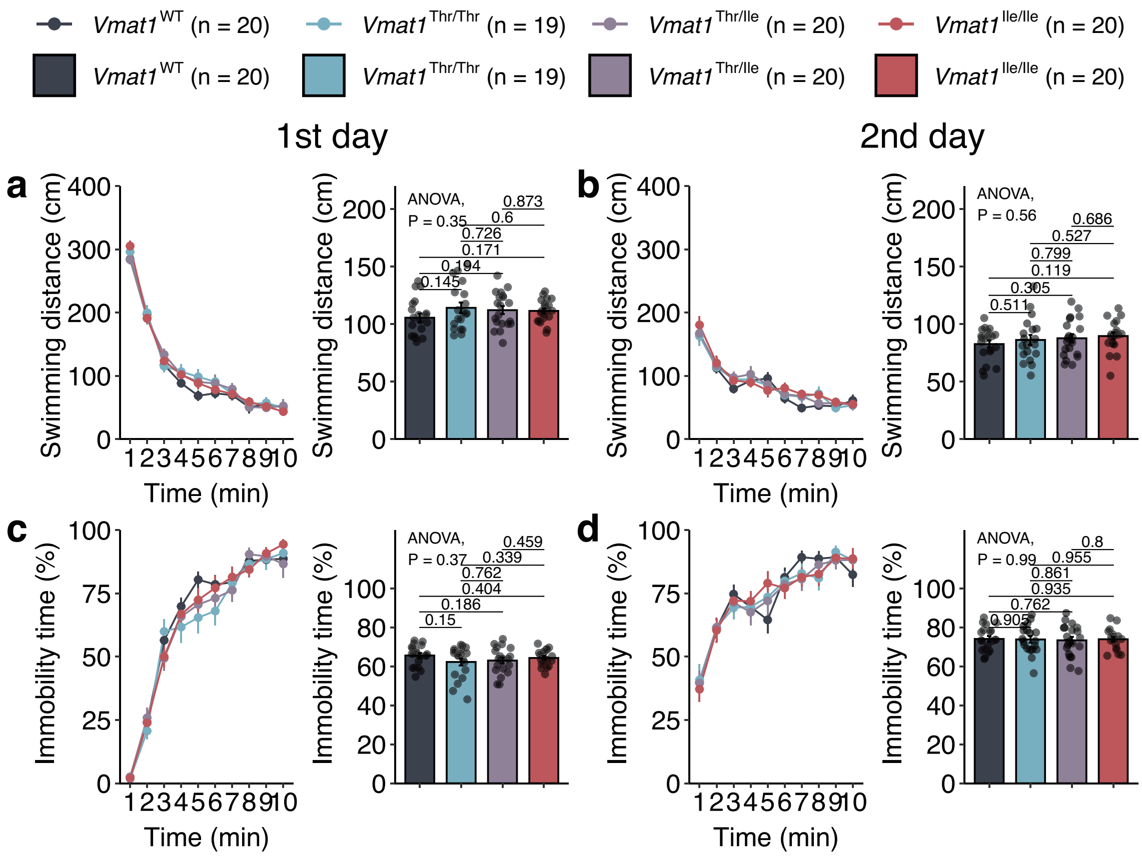


**Supplementary Figure S12. Porsolt forced swimming test results. (a)** Total swimming distance on the 1st day, **(b)** total swimming distance on the 2nd day, **(c)** immobility time within each time bin (% of total bin time) on the 1st day, and **(d)** immobility time within each time bin on the 2nd day. For each parameter, the left panel plots the value over time and the right panel plots the average for the entire trial. *P* values were calculated by one-way ANOVA and pair-wise *t*-tests (uncorrected). Error bars represent standard errors.

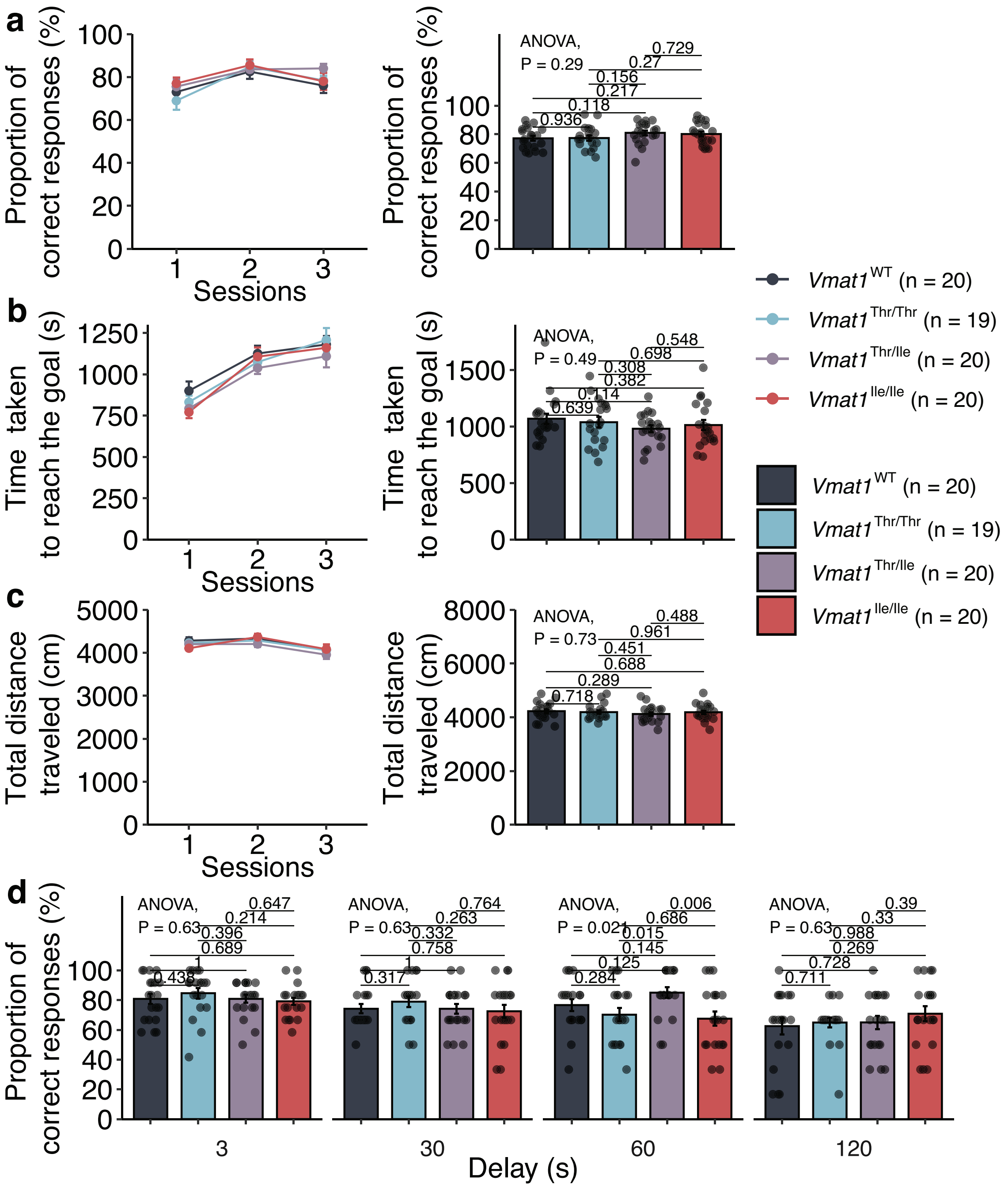


**Supplementary Figure S13. T-Maze test results. (a)** Proportion (%) of correct responses, **(b)** time taken to reach the goal, **(c)** total distance traveled, and **(d)** proportion (%) of correct responses in trials with delay. For each parameter shown in **(a)** to **(c)**, the left panel plots the value with time and the right panel plots the average over the entire trial. *P* values were calculated by one-way ANOVA and pair-wise *t*-tests (uncorrected). Error bars represent standard errors.

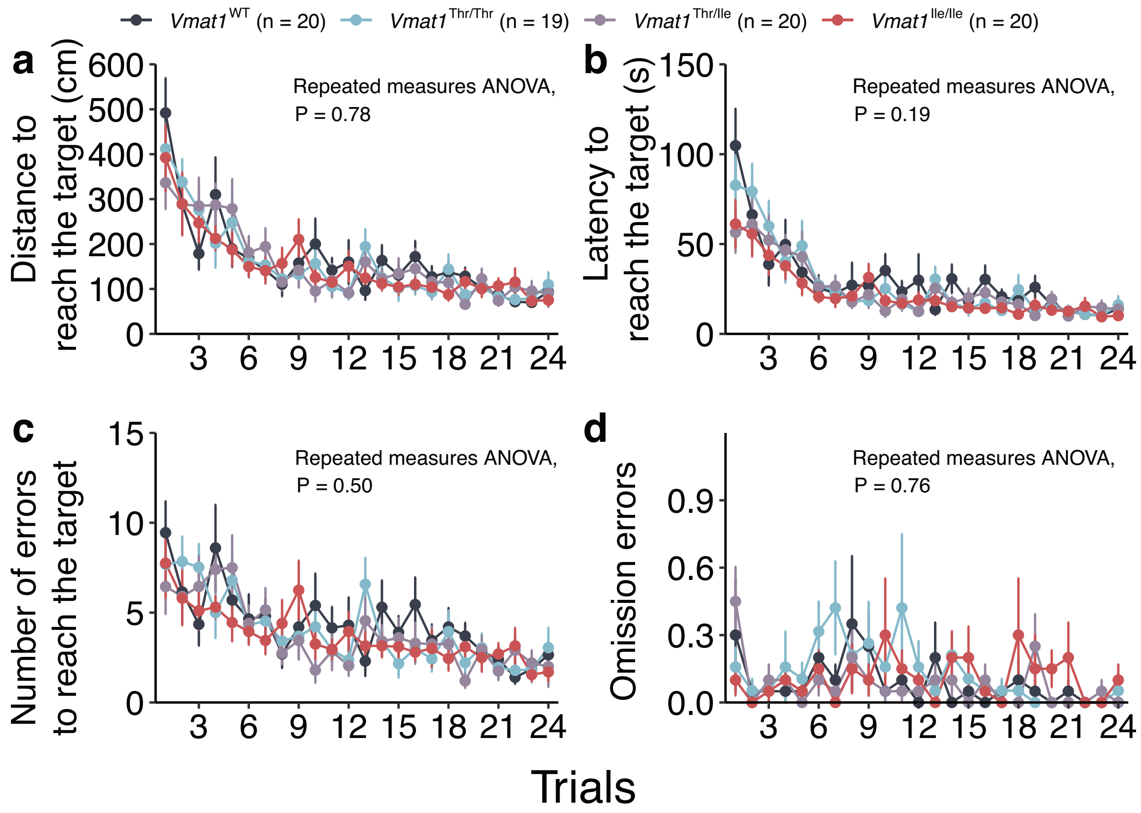


**Supplementary Figure S14-1. Barnes maze trial results. (a)** Average distance traveled, **(b)** latency, and **(c)** number of errors to reach the target hole on each trial. **(d)** Omission errors on each trial. *P* values were calculated by two-way repeated measures ANOVA. Error bars represent standard errors.

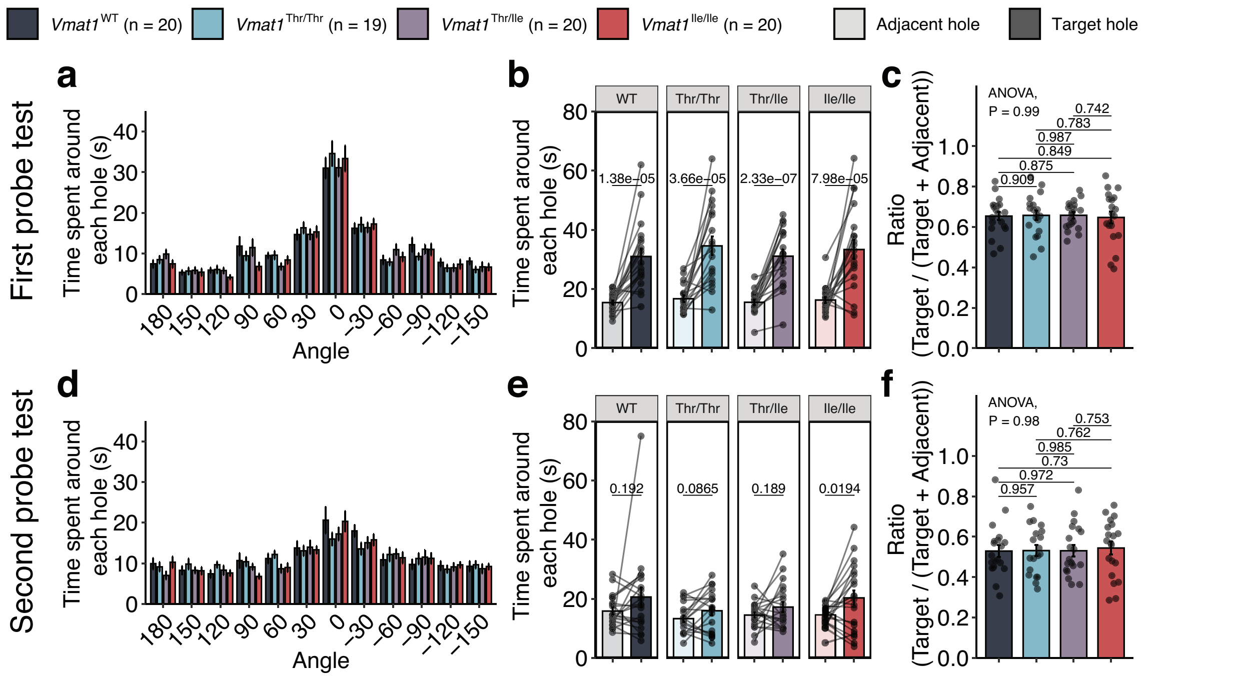


**Supplementary Figure S14-2. Barnes maze probe test results. (a–c)** First probe test results. **(d-f)** Second probe test results. **(a)** and **(d)** Time spent around each hole, where the angle “0” represents the target hole. **(b)** and **(e)** Time spent around the target and adjacent holes (30° and –30° holes). **(c)** and **(f)** Ratio of time spent around the target hole to total time spent around the target and adjacent holes. *P* values were calculated by paired *t*-tests in **(b)** and **(d)**, and by one-way ANOVA in **(c)** and **(e)**. Error bars represent standard errors.

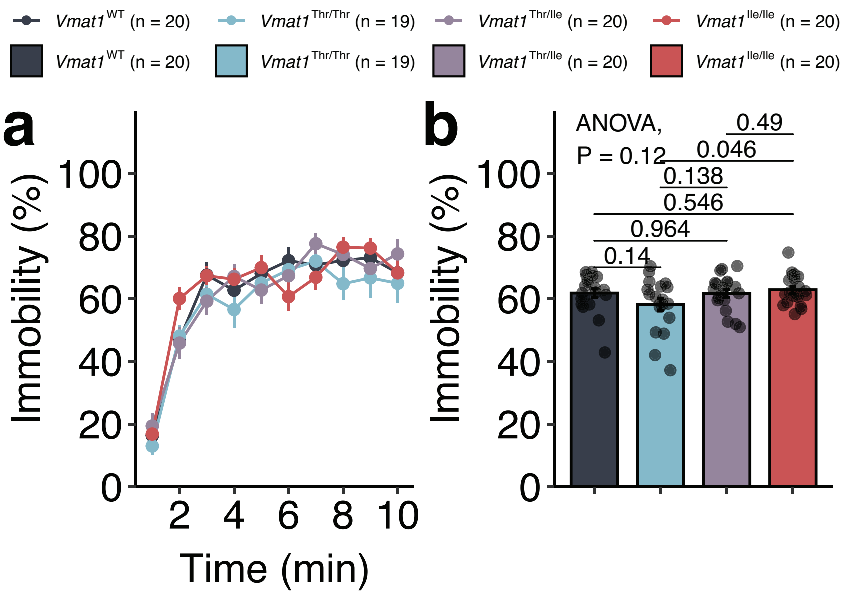


**Supplementary Figure S15. Immobility during the tail suspension test.** The left panel plots relative immobility time within successive time bins (% of bin duration), while the right panel plots the average relative immobility time over the entire test (% of total test duration). *P* values were calculated by one-way ANOVA and pair-wise *t*-tests (uncorrected). Error bars represent standard errors.

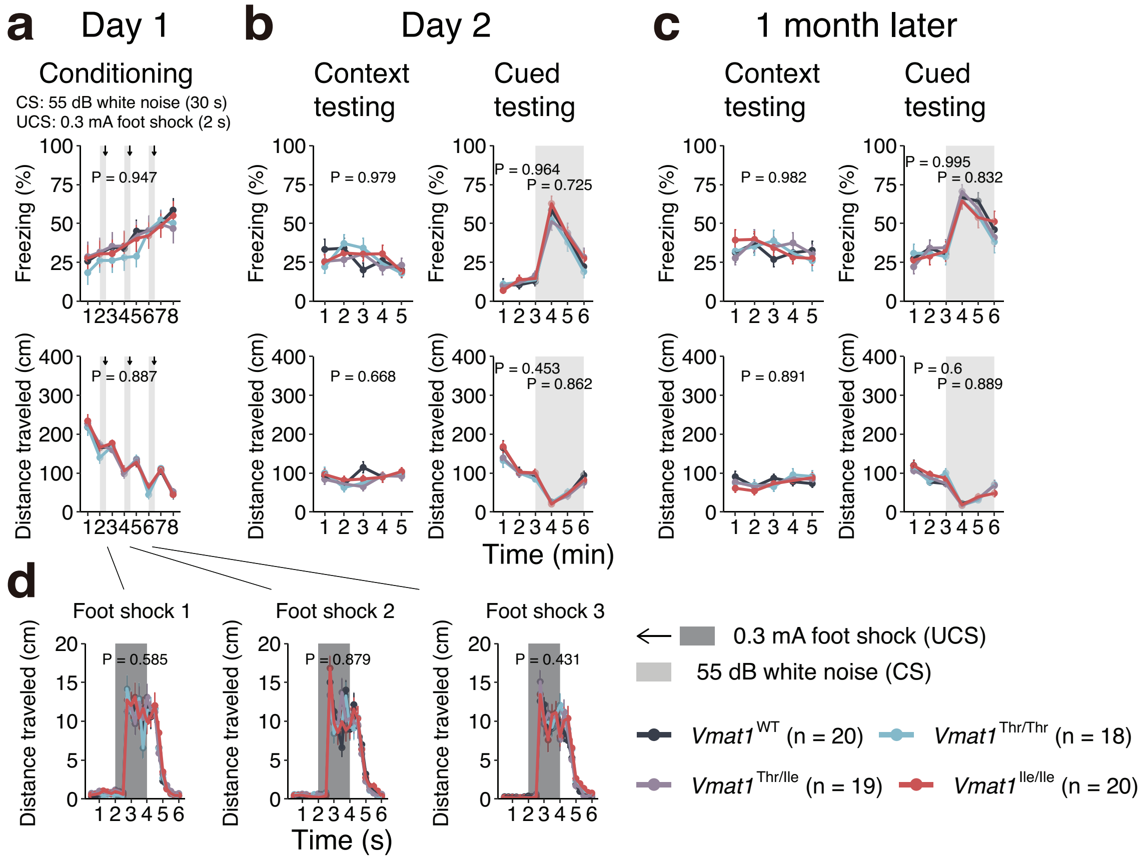


**Supplementary Figure S16. Fear conditioning test results. (a)** Freezing ratio and total distance traveled during the conditioning tests. Mice were exposed to a 55-dB white noise tone (CS) followed by 0.3-mA foot shocks (US) delivered at three times within 8 minutes. **(b)** and **(c)** Freezing ratio and distance traveled during the context and cued tests on the second day **(b)** and one month later **(c)**. **(d)** Distance traveled during the foot shock. Light gray and gray shadows represent CS and US, respectively. *P* values were calculated by two-way repeated measures ANOVA. Error bars represent standard errors.

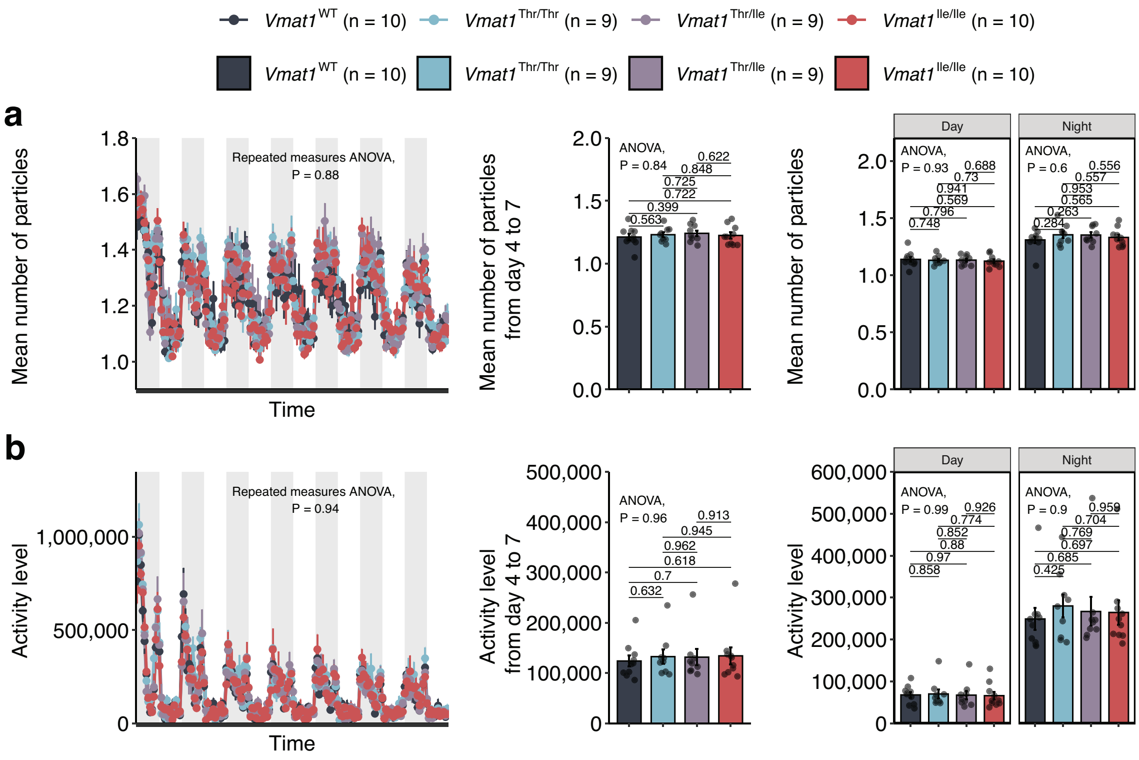


**Supplementary Figure S17. Home cage social interaction test. (a)** The mean number of particles (mouse tracer symbols) detected and **(b)** activity levels (in arbitrary units). The gray shadow indicates night (from 7:00 PM to 7:00 AM). The left panel plots the changes with time, the middle panel plots the average from day4 to 7, and the right panel plots the day average and night average throughout the entire test. *P* values were calculated by two-way repeated measures ANOVA, one-way ANOVA, and pair-wise *t*-tests (uncorrected). Error bars represent standard errors.

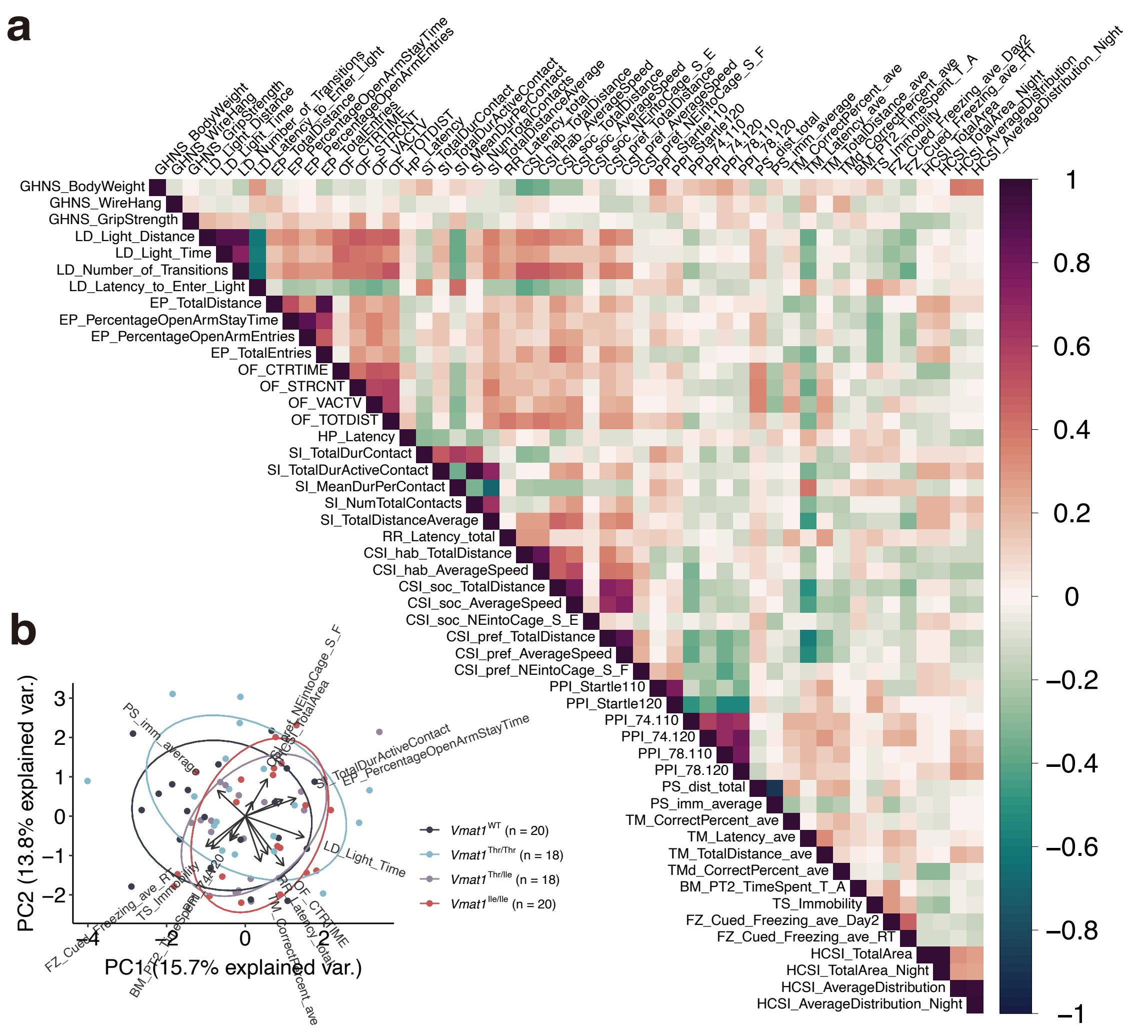


**Supplementary Figure S18. Overall behavioral patterns throughout the test battery. (a)** Pearson's correlations among test scores. **(b)** Principal component analysis was conducted to visualize the behavioral pattern of individual mice. A variable was selected as a representative from each test and used to calculate principal component scores. In both panels, only individuals with complete test scores (all tests) were included (n = 20, 18, 16, and 18 for *Vmat1*$^{WT}$, and *Vmat1*$^{Thr/Thr}$, *Vmat1*$^{Thr/Ile}$, and *Vmat1*$^{Ile/Ile}$, respectively). Pairs in SI and HCSI tests received the same score. In the wire hanging test, durations longer than 60 s were recorded as 60 s.

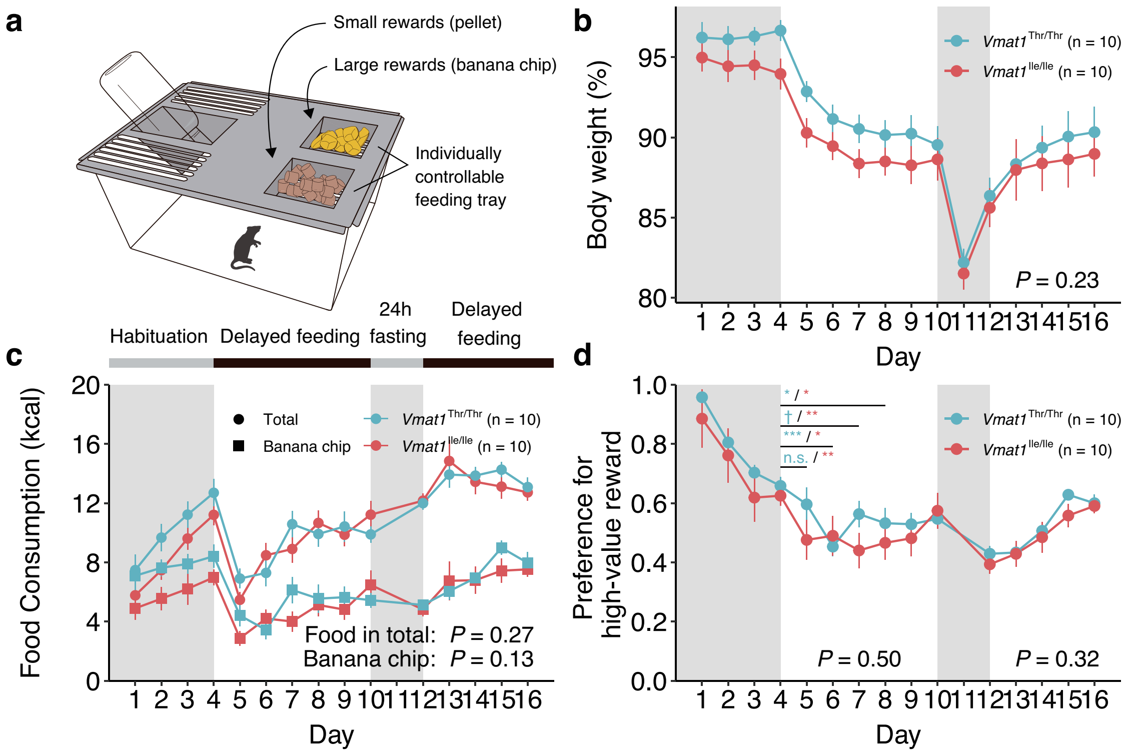


**Supplementary Figure S19. Impulsivity tests. (a)** Cage used for the impulsivity test. Two trays are individually controllable by a computer program. Time-series data of **(b)** body weight, **(c)** food consumption, and **(d)** preference for high-value reward (banana chip) on each day of the test. Blue and red point represents *Vmat1*$^{\mathrm{Thr}}$ and *Vmat1*$^{\mathrm{Ile}}$ genotype, respectively. In **(c)**, Rectangles and closed circles represent banana chips and total consumption of foods, respectively. Difference in values between genotypes was assessed by generalized additive model with gaussian distribution for **(b)** and **(c),** and with quasi-binomial distribution for **(d)** separately for day 4 to 10 (after habituation) and 12 to 16 (after fasting). Statistical significance was also evaluated by paired t-test in **(d)** to compare the preference to the basal level after habituation (Day4). †: $0.05 < P < 0.1$, *: $0.01 < P < 0.05$, **: $0.001 < P < 0.01$, ***: $P < 0.001$. Error bars represent standard errors.

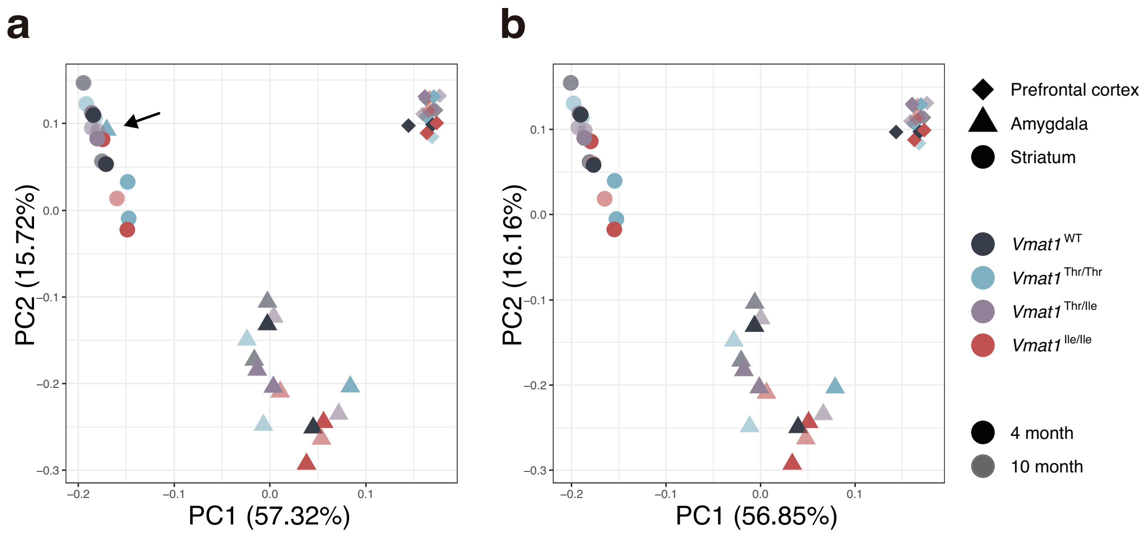


**Supplementary Figure S20. Principal component analysis of the gene expression profile for each brain sample. (a)** Principal component analysis of 48 samples and **(b)** 47 samples excluding a suspicious sample from the amygdala of a *Vmat1*$^{\mathrm{Thr/Thr}}$ mouse indicated by an arrow in **(a)**. The shape, color, and gray-scale intensity of the plot represent the brain region, genotype, and age, respectively.

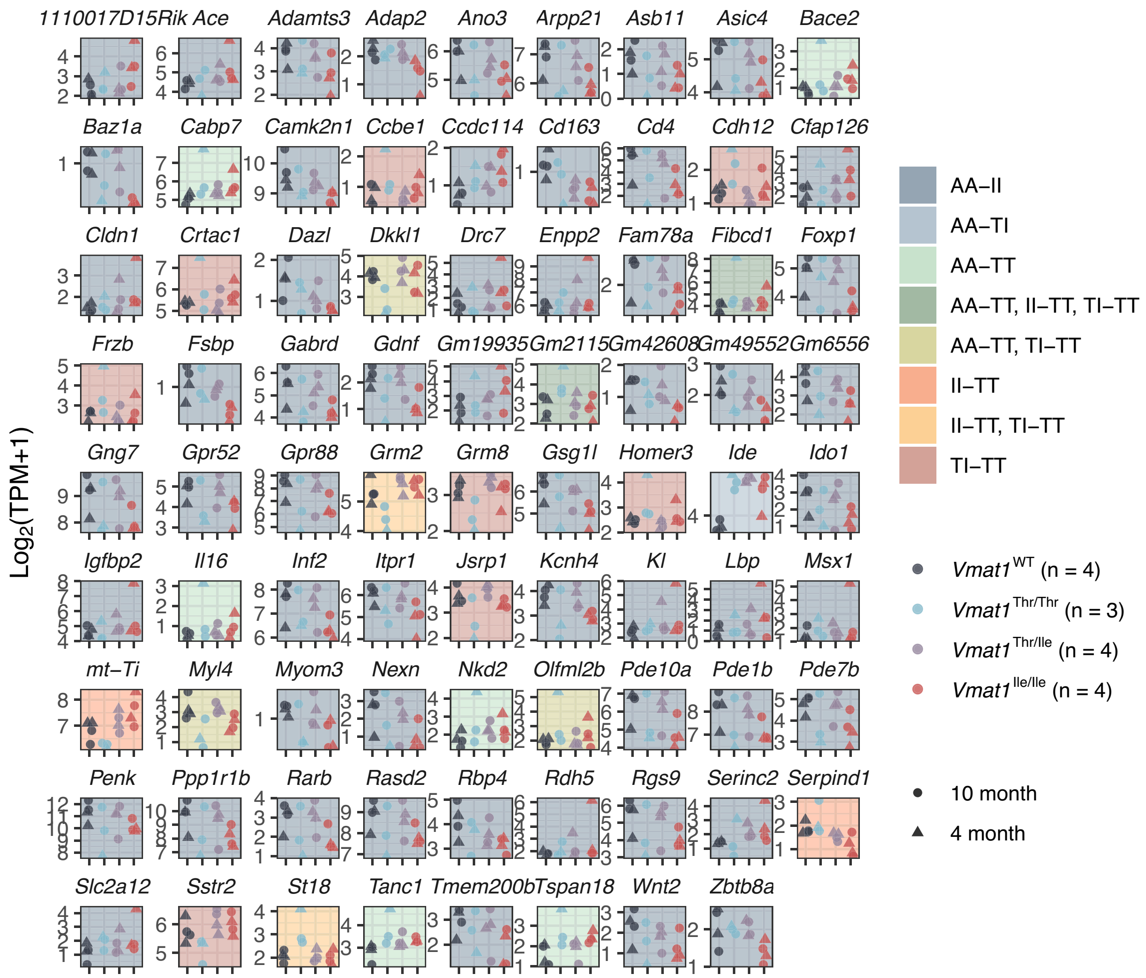


**Supplementary Figure S21. Expression levels of DEGs detected among *Vmat1* genotypes in the amygdala.** The shape and the color of the data points denote the age and genotype of the sample, respectively. Each panel is highlighted in a color corresponding to the pair-wise identifying DEG.

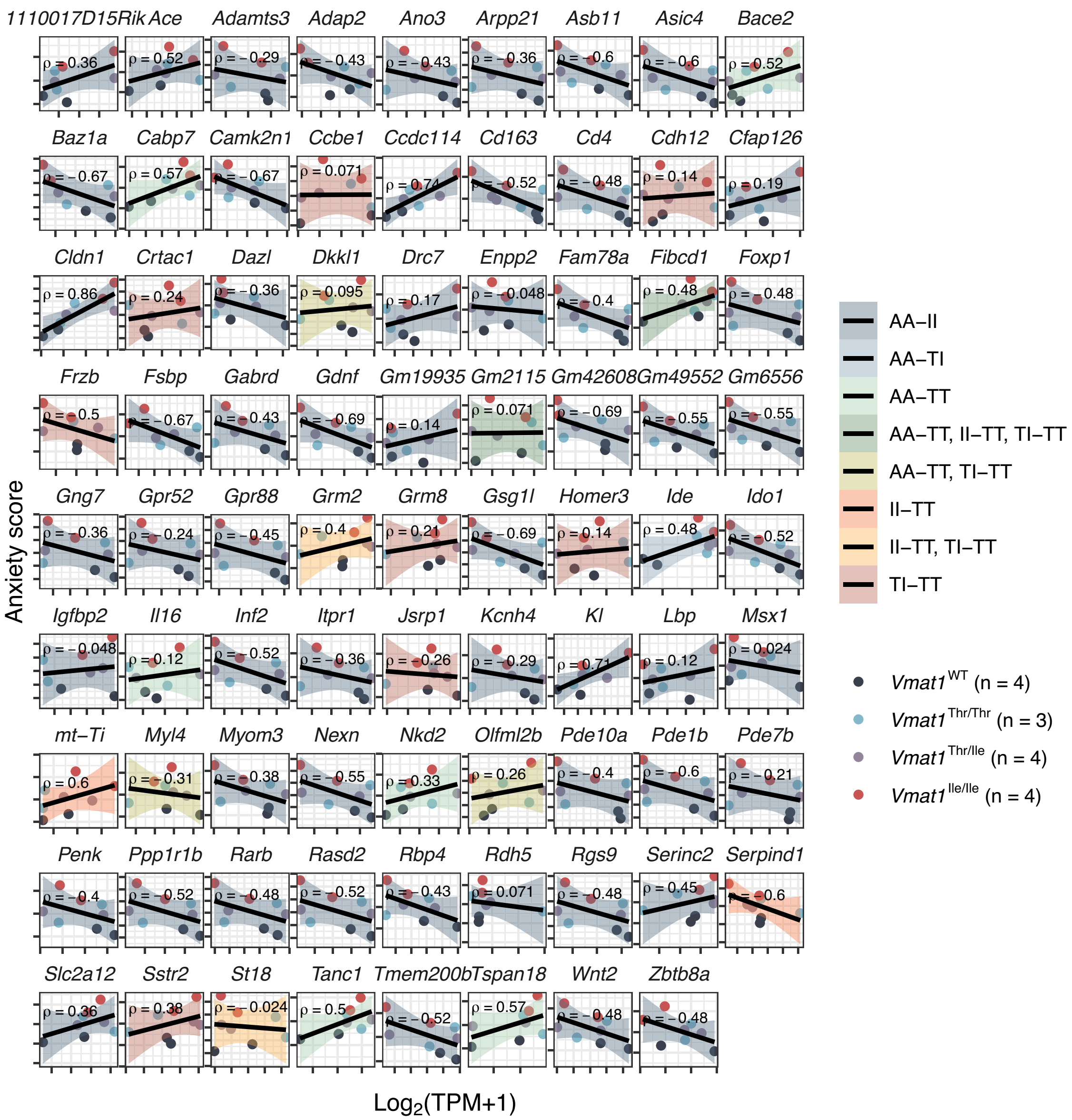


**Supplementary Figure S22. Correlations of individual expression levels of DEGs with anxiety scores.** In each panel, the linear regression line with Spearman's correlation coefficient is presented, and the color of the 95% confidence interval corresponds to the specific pair-wise comparison identifying the DEG.

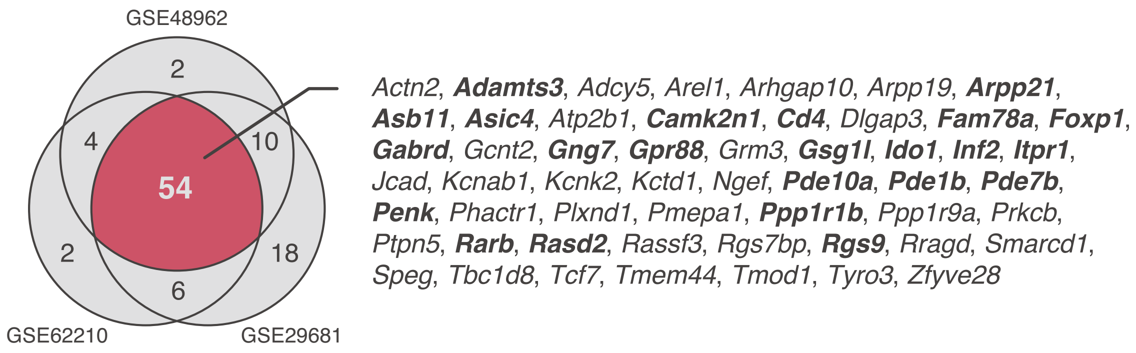


**Supplementary Figure S23. DEGs overlapped among three RNA-seq datasets of Huntington disease and the current dataset.** The DEGs detected in three previous studies showed the large overlap among each other (54 genes), out of which genes in bold on the right indicate the DEGs detected between WT vs. Ile comparison in the present study.

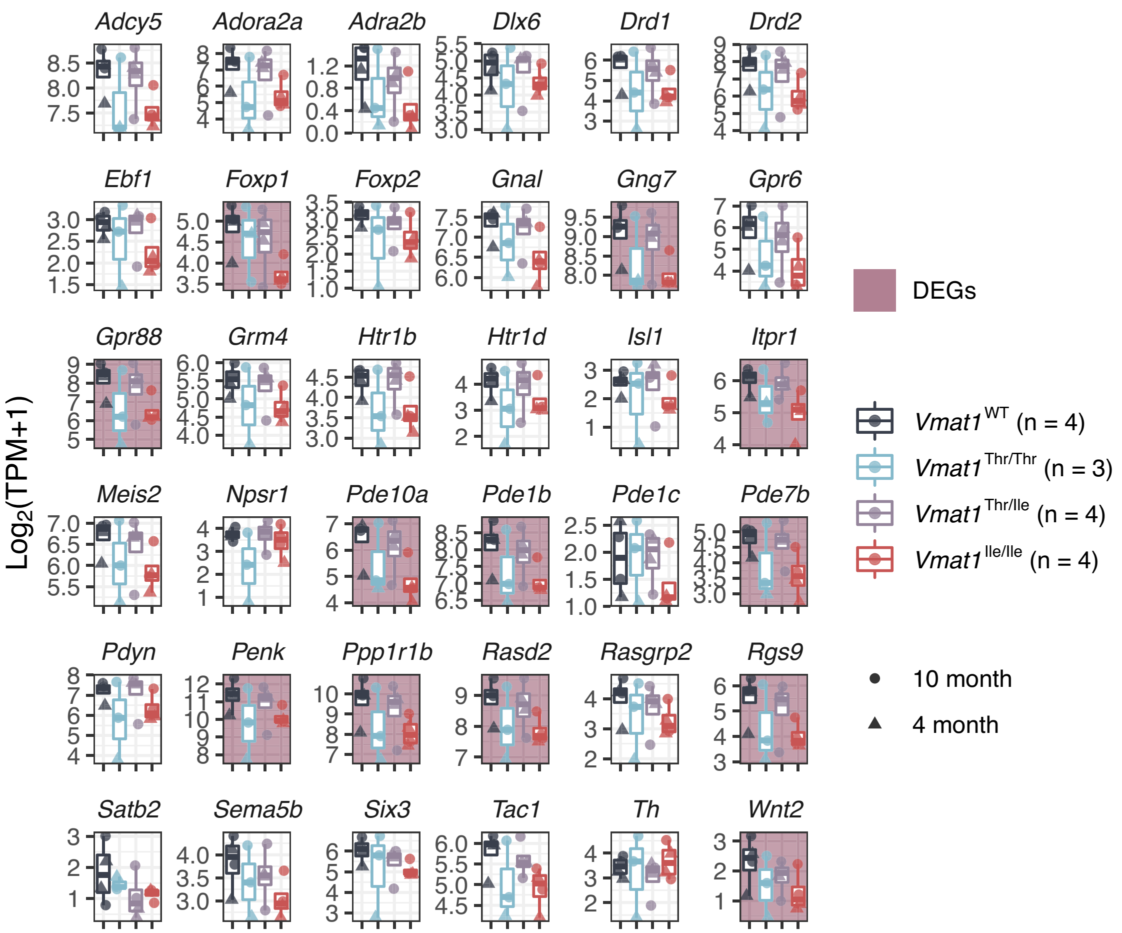


**Supplementary Figure S24. Expression levels of the genes involved in monoaminergic/neuropeptide signaling or neurogenesis and belonging to the same co-expressing module.** Genes highlighted in red are DEGs detected between *VMAT1*$^{\mathrm{WT}}$ and *VMAT1*$^{\mathrm{Ile/Ile}}$ mice. Note that *Th* is not included in the M1 module but is shown for comparison (see Discussion).

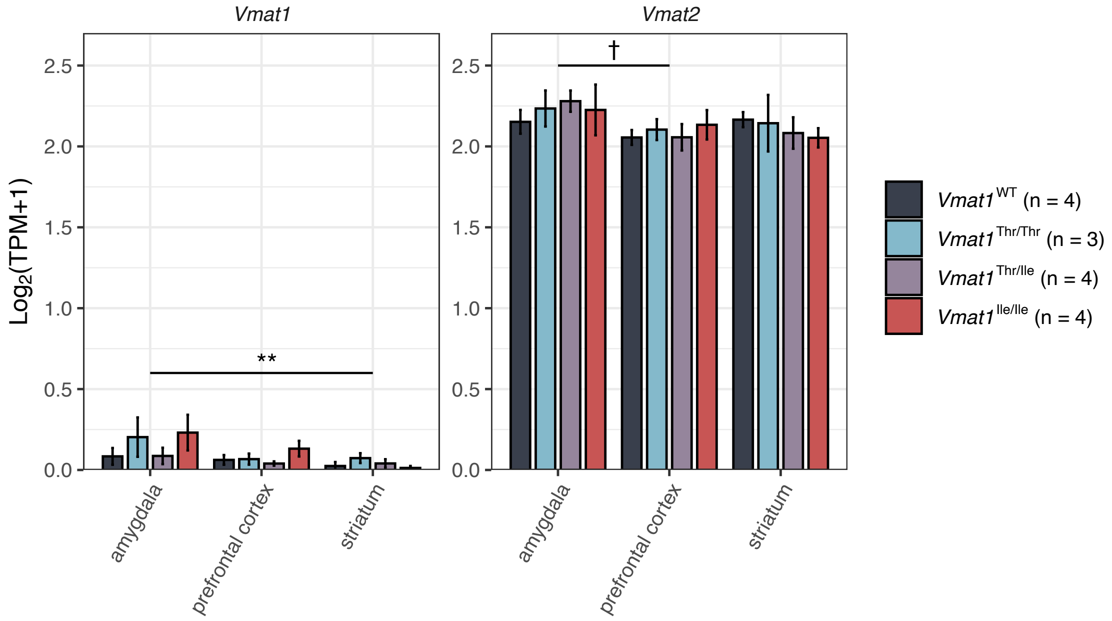


**Supplementary Figure S25. Expression levels of *Vmat1* and *Vmat2* in prefrontal cortex, amygdala, and striatum.** (Left) VMAT1 expression was higher in samples from the than samples from striatum (*P* = 0.0097). (Right) VMAT2 was expressed at comparable levels across regions with only a marginal difference (*P* = 0.071) between the amygdala and prefrontal cortex. Statistical significance was tested by Dunnett's test and samples were merged across genotypes. Error bars represent standard errors.
